# Phylogenomic analysis of *Clostridium perfringens* identifies isogenic strains in gastroenteritis outbreaks, and novel virulence-related features

**DOI:** 10.1101/670448

**Authors:** Raymond Kiu, Shabhonam Caim, Derek Pickard, Anais Painset, Craig Swift, Gordon Dougan, Alison E Mather, Corinne Amar, Lindsay J Hall

**Author notes:** **Corresponding author**, Lindsay J Hall.

## Abstract

*Clostridium perfringens* is a major enteric pathogen known to cause gastroenteritis in human adults. Although major outbreak cases are frequently reported, limited Whole Genome Sequencing (WGS) based studies have been performed to understand the genomic epidemiology and virulence gene content of *C. perfringens*-associated outbreak strains. We performed both genomic and phylogenetic analysis on 109 *C. perfringens* strains (human and food) isolated from disease cases in England and Wales between 2011-2017. Initial findings highlighted the enhanced discriminatory power of WGS in profiling outbreak *C. perfringens* strains, when compared to the current Public Health England referencing laboratory technique of Fluorescent Amplified Fragment Length Polymorphism (fAFLP). Further analysis identified that isogenic *C. perfringens* strains were associated with nine distinct care home-associated outbreaks over the course of a 5-year interval, indicating a potential common source linked to these outbreaks or transmission over time and space. As expected the enterotoxin CPE gene was encoded in all but 4 isolates (96.4%; 105/109), with virulence plasmids encoding *cpe* (particularly pCPF5603- and pCPF4969-family plasmids) extensively distributed (82.6%;90/109). Genes encoding accessory virulence factors, such as beta-2 toxin, were commonly detected (46.7%; 50/109), and genes encoding phage proteins were also frequently identified, with additional analysis indicating their contribution to increased virulence determinants within the genomes of gastroenteritis-associated *C. perfringens*. Overall this large-scale genomic study of gastroenteritis-associated *C. perfringens* suggested that 3 major sub-types underlie these outbreaks: strains carrying (1) pCPF5603 plasmid, (2) pCPF4969 plasmid, and (3) strains carrying *cpe* on transposable element Tn*5565*(usually integrated into chromosome). Our findings indicate that further studies will be required to fully probe this enteric pathogen, particularly in relation to developing intervention and prevention strategies to reduce food poisoning disease burden in vulnerable patients, such as the elderly.

## Introduction

*Clostridium perfringens* is an important pathogen known to cause disease in humans and animals (1, 2). Notably, the pathogenesis of *C. perfringens*-associated infections is largely attributed to the wide array of toxins this species can produce, with >20 toxins currently identified (3, 4). This Gram-positive spore former has been associated with foodborne and non-foodborne diarrhoeal diseases in humans, and preterm-necrotising enterocolitis (5, 6).

*C. perfringens-associated* food poisoning, also termed acute watery diarrhoea, was first documented in the UK and USA in the 1940s (7). Typical symptoms occur within 8-14 h after ingestion of food contaminated with at least 10^6^ CFU/g of live bacterial cells. These include intestinal cramp, watery diarrhoea without fever or vomiting, which normally resolves in 12-24 h (8). Importantly, *C. perfringens* is currently the second most common foodborne pathogen in the UK after *Campylobacter*, with cases often under reported due to the frequently self-limiting nature of the illness, with current estimates suggesting ~80,000 cases/annum (9–12).

In the UK, antibiotic-associated diarrhoea and non-foodborne outbreaks of *C. perfringens* diarrhoea have been frequently reported since the 1980s amongst the elderly, particularly in hospital settings (13). With this type of illness, symptoms are more severe than foodborne diarrhoea and are longer lasting (>3 days to several weeks), often chronic, and infections are more likely to be spread amongst cases (14). This type of *C. perfringens* infection has also been reported in elderly patients, especially those residing in care homes in the North East of England between 2012-2014 (83% of the outbreaks reported from care homes) (10). Although fatality due to *C. perfringens* diarrhoea is uncommon and hospitalisation rate is low, enterotoxigenic *C. perfringens* is reported to cause ~55 deaths/year in England and Wales according to the UK Food Standards Agency (15, 16).

The newly expanded and revised toxinotyping scheme classifies *C. perfringens* into 7 toxinotypes (type A-G) according to the combination of typing toxins produced, with this classification used in this article (17). Human cases of *C. perfringens* diarrhoea are primarily caused by type F strains (formerly classified as enterotoxigenic type A), which produce enterotoxin (CPE), encoded by the *cpe* gene (18). This potent pore-forming toxin is reported to disrupt intestinal tight junction barriers, which is associated with intestinal disease symptoms (19). *C. perfringens*, and associated encoded toxins, have been extensively studied with respect to disease pathogenesis, with a strong focus on animal infections (20–24). Recent studies analysing a range of diverse *C. perfringens* strains (from both animal and human-associated infections) indicates a plastic and divergent pangenome, with a significant proportion of accessory genes predicted to be involved in virulence mechanisms and metabolisms, linked to enhanced host colonisation and disease initiation (3, 25). However, studies describing human outbreak-associated *C. perfringens* infections are limited, and to date only one recent study (58 isolates) has utilised Whole Genome Sequencing (WGS) data to probe the genomic epidemiology of associated strains (3, 26).

We have applied in-depth genomics and phylogenetic analyses to whole genome sequences of 109 newly-sequenced *C. perfringens* isolates associated with outbreaks or incidents of *C. perfringens* diarrhoea in England and Wales, either foodborne or non-foodborne-derived. We also identified distribution of known virulence-related determinants including toxin and antimicrobial resistance (AMR) genes, virulence-associated plasmid contents within food and case isolates and probed putative functional capabilities of the accessory genomes and virulence features within encoded phage genomes. Importantly, we determined that isogenic strains were associated with 9 care-home outbreaks in North East England between 2013-2017, and furthermore uncovered the significant involvement of virulence plasmid-carrying *C. perfringens* in these outbreaks.

## Materials and methods

### Bacterial isolates, PCR and genomic DNA extraction

*C. perfringens* isolated from clinical cases of diarrhoea and suspected foods when available, were referred to the reference laboratory at PHE, Gastrointestinal Bacteria Reference Unit (GBRU). Identification and characterisation of cultures was performed by detection of the *C. perfringens* alpha toxin and enterotoxin gene by duplex real time PCR as described elsewhere (27). Enterotoxigenic *C. perfringens*, when associated with an outbreak or incident, were then further typed for strain discrimination using fluorescent amplified fragment length polymorphism (fAFLP) as previously described (28). In this study, 109 cultures characterised and archived by the GBRU between 2011 and 2017 were selected, representing enterotoxigenic and non-enterotoxigenic isolates from sporadic cases and outbreaks of *C. perfringens* food poisoning and of non-foodborne *C. perfringens* diarrhoea (**Table S1**).

DNA was extracted from overnight *C. perfringens* cultures (maximum of 1 x 10^7^cells) lysed with lysis buffer (200μl Qiagen Buffer P1, 20 μl lysozyme with concentration at 100mg/ml) and incubated at 37°C for 1 h before addition of 20 μl Proteinase K (QIAsymphony DSP DNA kit), and incubation at 56°C for 5 h until cells had visibly lysed. Following incubation at 96°C for 10 min, to inactivate proteases and any viable cells remaining, RNA was removed by adding 0.4 mg of RNAse (Qiagen, Manchester) and incubation at 37°C for 15 min.

Genomic DNA was then purified using the QIAsymphony DSP DNA kit on a QIAsymphony SP automated DNA extraction platform (Qiagen, Manchester) according to manufacturer instructions.

### Computing infrastructure

Computational analyses were performed on Norwich Biosciences Institute’s (NBI) High Performing Computing cluster running Linux CentOS v7. Open-source software was utilised for genomic analysis.

### Genomic DNA sequencing

Pure bacterial DNA was subjected to standard Illumina library preparation protocol prior to sequencing on in-house Illumina MiSeq (PH091-PH156) or HiSeq 2500 platforms (PH004-PH090; at Wellcome Trust Sanger Institute, UK) with read length (paired-end reads) 2 x 101 bp and 2 x 151 bp respectively.

### Genome assembly and annotation

Sequencing reads were assembled using SPAdes v3.11 (PH091-PH156) to generate draft genomes, the remaining assemblies were generated at Wellcome Trust Sanger Institute (Hinxton, UK) as described previously (for assembly quality see **Table S2**)(29, 30). Assemblies were improved by scaffolding/gap filling using SSPACE v3.0 and GapFiller v1.10 (31–33). Sequencing reads and coverage counts were calculated via in-house custom script using FASTQ reads and FASTA assemblies. Genome assemblies were annotated by Prokka v1.13 using in-house Genus-specific database that included 35 *Clostridium* species retrieved from NCBI RefSeq database to construct genus-specific annotation database (**Table S3**)(34). Sequences from very small contigs (contig size <200 bp) were removed prior to coding region prediction. Assembly statistics were extracted from Prokka outputs using an in-house script. Draft genomes were checked for sequence contamination using Kraken v1.1 (based on MiniKraken database) (35). All draft genomes containing >5% of contaminated species sequences (other than *C. perfringens)*, whole genome Average Nucleotide Identity (ANI) < 95% (compared with reference genome NCTC8239; performed with Python module pyani v0.2.4), and assemblies with >500 contigs (considered as poor assemblies) were all excluded from further study (n=21)(36).

### Reference genome

A newly sequenced historical foodborne isolate genome NCTC8239 under the NCTC3000 project^1^ was retrieved (accession: SAMEA4063013) and assembled in this study with Canu pipeline v1.6 using PacBio reads (37). The final high-quality assembled genome consists of 3 008 497 bp in 2 contigs (contig 1 has 2 940 812 bp, contig 2 has 67 685 bp).

### Pangenome and phylogenetic analyses

Pangenome of isolates was constructed using Roary v3.8.0 at BLASTp 90% identity, adding option -s (do not split paralogs), and options -e and -n to generate core gene alignment using MAFFT v7.3 (38, 39). Roary took GFF3-format annotated assemblies generated by Prokka. The pangenome includes both the core and accessory genomes; core genome is defined as genes present in at least 99% of the genomes, accessory genome as genes present in <99% of the genomes. SNP-sites v2.3.3 was used to extract single nucleotide variants from the core gene alignment for re-constructing a phylogenetic tree (40). Phylogenetic trees were generated using FastTree v2.1.9 and annotated using iTOL v4.2 (41, 42). FastTree was run using the Generalized Time-Reversible (GTR) model of nucleotide evolution on 1000 bootstrap replicates to generate maximum-likelihood trees (41). SNP distance between genomes were computed using snp-dists (43). Population structure was analysed via the Bayesian-based clustering algorithm hierBAPS to assign lineages, implemented in R using *rhierbaps* v1.0.1 (44, 45). The pangenome was visualised in Phandango while plots were generated using the associated R scripts in the Roary package (46).

### Profiling virulence and plasmid-related sequences

Screening of toxin and AMR gene profiles, IS elements and plasmid *tcp* loci were performed via ABRicate with 90% identity and 90% coverage minimum cutoffs to infer identical genes based on a custom toxin database and the CARD database v2.0.0 (AMR) as described previously (47, 48). ARIBA v2.8.1 was used as a secondary approach to confirm detections of both toxin and AMR genes in raw sequence FASTQ files (49). Heat maps were generated in R using *gplots* and function *heatmap.2* (50, 51).

### *In silico* plasmid analysis

Sequencing reads were utilised for computational plasmid prediction via software PlasmidSeeker v1.0 (52). Plasmid prediction was based on 8 514 plasmid sequences available in NCBI Reference Sequence databases (RefSeq; including 35 *C. perfringens* plasmids, see **Table S4**). All reads were searched for matching k-mers at k-mer length of 20 and screening cutoff at P-value 0.05 based on FASTQ reads. The top predicted plasmids appearing in each ‘cluster’ (with highest k-mer identity; k-mer percentage ≥80% as the minimum cutoff) were extracted as predicted plasmids (**Table S5**). Binary heat maps were generated as described in the previous section. Plasmid sequences from high-sequencing-coverage assemblies (>200X; single contig; n=12) were extracted using in-house Perl scripts, identified by plasmid gene content. Plasmid annotation was performed via Prokka v1.13 using *Clostridium* genus-specific database as described in the previous section. Plasmid comparison and visualisation were performed using Easyfig v2.2.2.

### Bacterial genome-wide association analysis

To associate subsets of genes with specific outbreaks or isolates, we used Scoary v1.6 to identify statistically-related genes based on Roary output (53). Cutoffs were set as ≥80% sensitivity and 100% specificity. Specifically, for a Care Home (CH) and Food Poisoning (FP) subset comparison, sensitivity cutoff was set at ≥50% and specificity at 100%.

### Pangenome-wide functional annotation

Functional categories (COG category) were assigned to genes for biological interpretation via eggNOG-mapper v0.99.3 based on the EggNog database (bacteria) (54, 55). Genes of interest were extracted via in-house Perl scripts from Roary-generated pangenome references. Annotations and COG categories were parsed with in-house scripts. Bar plots were generated using GraphPad Prism v6.

### Prophage mining

Web tool PHASTER was utilised for detection of intact prophage existing in bacterial genomes (**Table S6**). Annotated GenBank files were submitted manually to the PHASTER web server and annotated data parsed with in-house scripts. The detection of phage was based on the scoring method and classification as described previously (56). Only intact phage regions within the genomes of completeness score >100 (of maximum 150) were analysed further (extracted by default using PHASTER web tool), annotated and colour-coded using Prokka v1.13 and Artemis for visualisation in EasyFig v2.2.2 and R package *gplots* function *heatmap.2* (57, 58). Bar plots were generated using GraphPad Prism v6.

### Nucleotide sequence accession numbers

Sequence data in this study will be submitted to the European Nucleotide Archive (ENA) and made available under accession PRJEB25764 upon acceptance of manuscript.

## Results

### Whole-genome based phylogenetic analysis reveals potential epidemiological clusters

Initially we analysed the population structure of all strains sequenced. We defined general food poisoning isolates as Food Poisoning (FP, n =74), and care home specific isolates as Care Home (CH, n =35) (**Fig. 1A-B**). Quality of the genomic assemblies of draft genomes was also determined (**Fig. 1C; Table S2**), with >70% of the isolate assemblies <200 contigs.

**Fig. 1.**
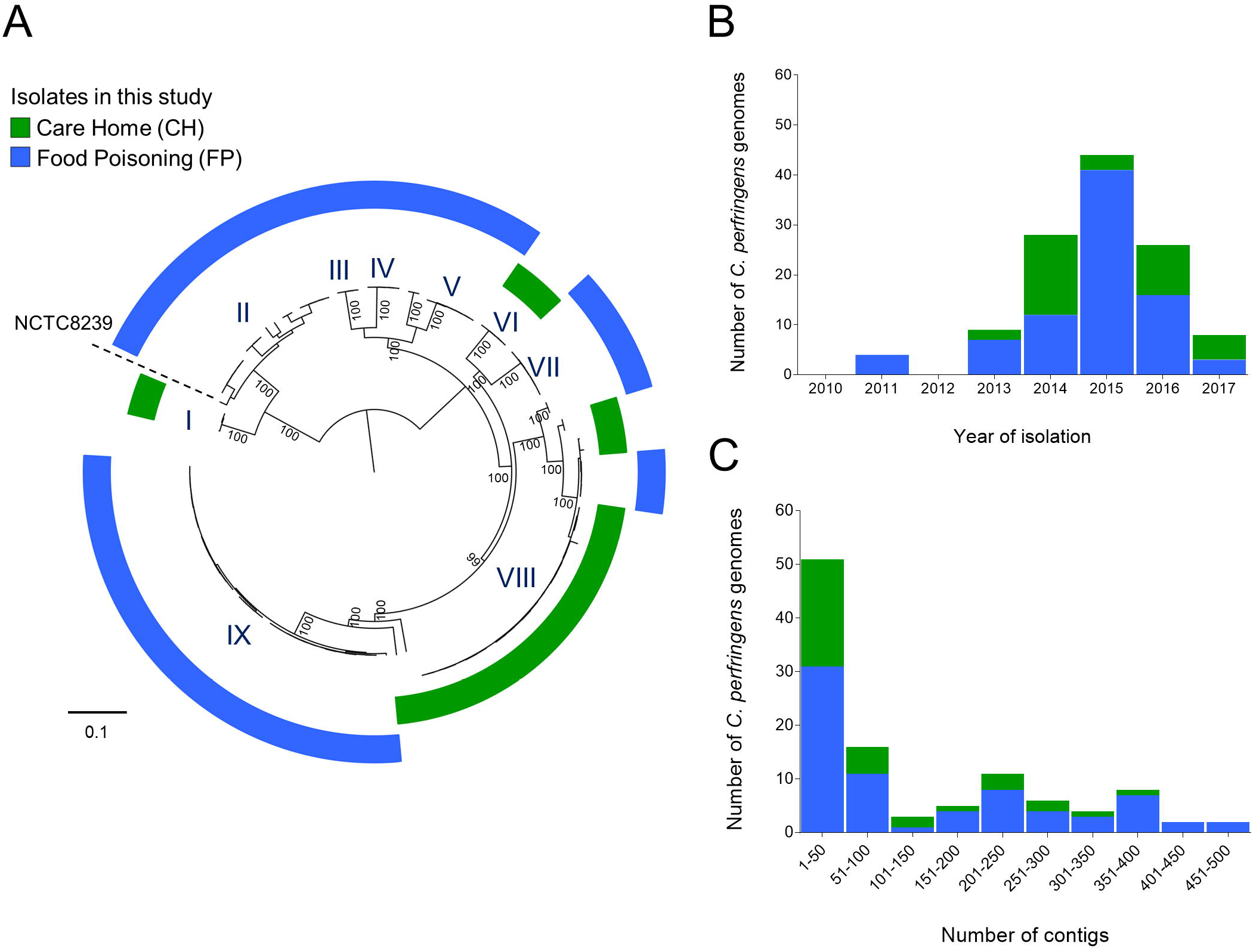
Population structure and sample distribution statistics for genome assemblies. **(A)** Population structure of 109 *C. perfringens* isolates analysed in this study. Mid-point rooted maximum-likelihood phylogeny inferred from 73 882 SNPs identified in 110 diarrhoea-associated *C. perfringens* isolates (including NCTC8239). The colour-coded rings indicated cohort-specific origins of isolates. Cluster VIII (green ring; clusters determined via hierBAPS clustering analysis) consists of primarily isolates obtained from multiple care home-associated outbreaks. Historical food poisoning isolate NCTC8239 was used as a public reference genome as indicated in the figure. Bootstrap values are represented in the tree. Branch lengths are indicative of the estimated nucleotide substitution per site (SNPs). **(B)** Temporal distribution of all 109 *C. perfringens* genomes included in this study. **(C)** Contig count distribution of *C. perfringens* genome assemblies in this study. More than 70% of the total assemblies are <200 contigs.

Separate analysis of CH isolates indicated four distinct phylogenetic lineages relating to care home outbreaks (**Fig. 2A**). Lineage I contained the reference genome NCTC8239, a historical *cpe*-positive isolate (originally isolated from salt beef) associated with a FP outbreak, and three newly sequenced strains (7). The remaining isolates clustered within three lineages (i.e. II, III and IV), that were divergent from lineage I indicating these CH isolates might be genetically distinct from typical FP isolates as in **Fig. 1A**. Further analysis indicated that 18 closely-related strains obtained from 9 different outbreaks between 2013-2017 which occurred in the North East England, clustered within the same IVc sub-lineage (**Fig. 2B**). SNP investigation on these IVc isolates determined within-sub-lineage pairwise genetic distances of <80 SNPs (29.9 ± 16.6 SNPs; mean ± S.D.; **Table S7**), suggesting a close epidemiological link. Isolates associated with specific outbreaks within sub-lineage IVc (i.e. outbreaks 2, 7, 8, 9 and 10) showed very narrow pairwise genetic distances <20 SNPs (6.6 ± 6.6 SNPs; mean ± S.D.; **Fig. 2C**), suggesting potential involvement of an isogenic strain (genetically highly similar) within these individual care home outbreaks (although a number of genetically dissimilar strains were also isolated from outbreaks 1, 2, 3, 6, 7 and 8 as shown in **Fig. 2A**).

**Fig. 2.**
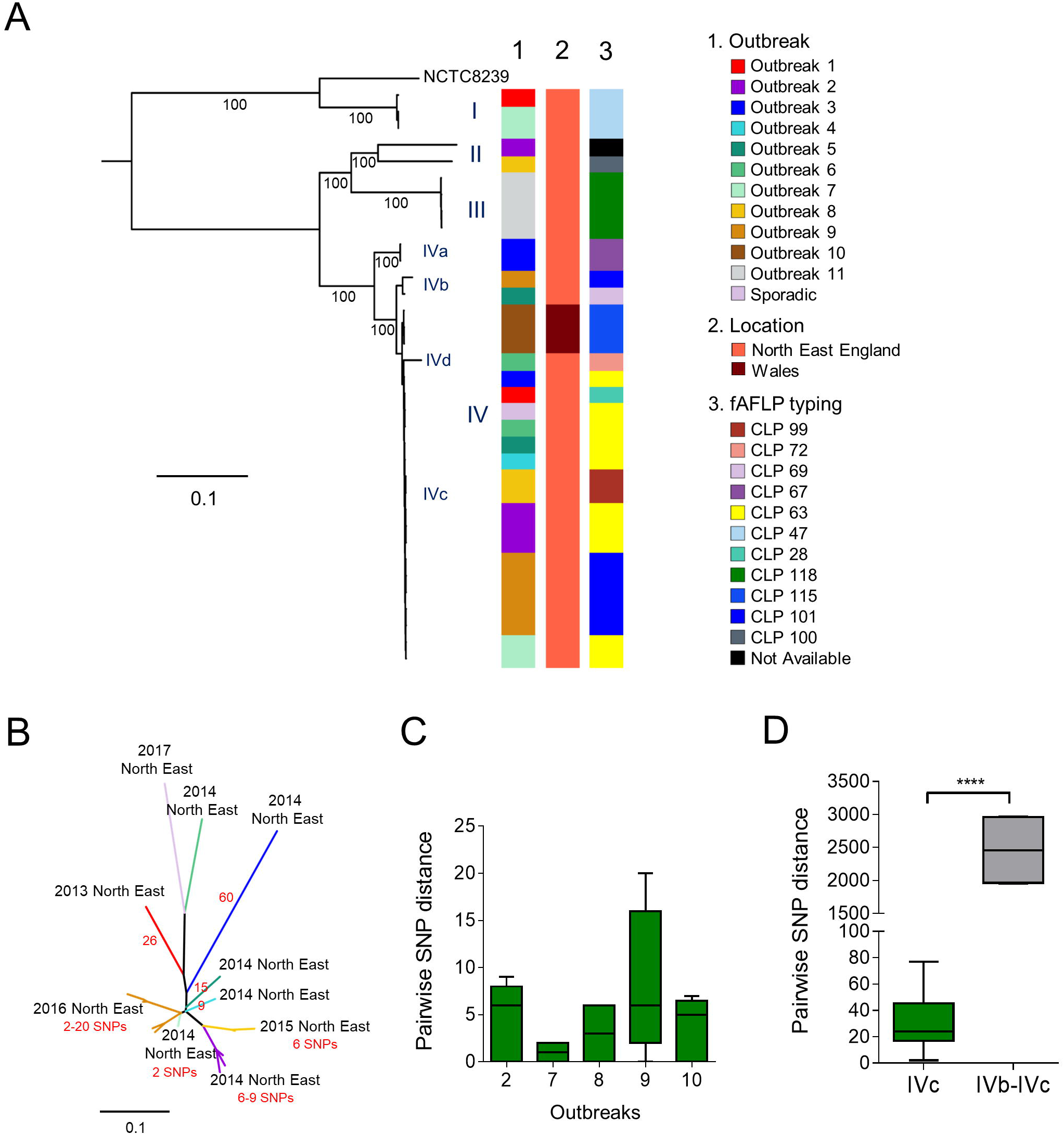
Phylogenomic analysis of care home-associated *C. perfringens* isolates. **(A)** Mid-point rooted maximum-likelihood phylogeny inferred from 64 560 SNPs (in core gene alignment) identified in 35 care home-associated *C. perfringens* isolates. The colour strips indicate diarrhoea outbreaks, location of outbreaks and fAFLP types respectively corresponding to the isolates. Branch lengths are indicative of the estimated SNP distance. Lineages and sub-lineages were determined via hierBAPS (level 1 & 2) clustering analysis. NCTC8239, a food poisoning isolate, was used as a public reference genome in this tree. Bootstrapping values are represented on the tree. **(B)** Unrooted maximum-likelihood tree (inferred from 191 SNPs in 18 genomes) of a sub-lineage IVc (excluding three genetically distant Welsh isolates) showing SNP distances in between 18 North-East England-derived isolates of individual outbreaks (labeled in locations and years, and SNP range in outbreaks; branches are colour-coded corresponding to individual outbreaks). SNP distances between branches are indicated in red numbers. **(C)** Pairwise within-outbreak core-SNP distance between isolates. **(D)** Pairwise outside-sub-lineage (IVb vs IVc) SNP comparison between isolates. Data: Mann-Whitney test. **** P<0.0001.

This WGS analysis was also shown to have greater discriminatory power than the currently used fAFLP. The fAFLP typing (type CLP 63, yellow-coded) failed to discriminate isolates from 6 different outbreaks (CH outbreaks 2-7; **Fig. 2A**), while SNP analysis clearly distinguished these strains (**Fig. 2B**) (59).

Analysis of FP isolates indicated clear separation between linages (**Fig. 3A**), particularly between lineage I, and remaining lineages II-VII (pairwise mean SNP distance lineage I vs lineages II-VII: 35165 ± 492 SNPs; within lineage I: 5684 ± 2498 SNPs; within lineages II-VII: 13542 ± 8675 SNPs). Isolates from three individual foodborne-outbreaks within lineage VII appear to be highly similar even though these isolates demonstrated geographical heterogeneity (**Fig. 3A**), and further analysis indicated two different outbreaks that occurred in London (2013) were related, but somewhat distinct from isolates obtained in North East England (2015) outbreaks (**Fig. 3B**). This suggests a geographical separation of a common ancestor at an earlier time point and may also indicate the potential widespread distribution of a genetically-related strain.

Isolates from individual FP outbreaks also appeared to be clonal and isogenic, as pairwise genetic distances were between 0-21 SNPs (mean genetic distance: 2.6 ± 2.7 SNPs; **Fig. 3C and Table S8**), when compared to same-lineage-between-outbreaks SNP distances of >1200 SNPs (**Fig. 3D**). In addition, outbreak-associated food source isolates were not distinguishable from human clinical isolates (genetically similar, pair-wise SNP range: 0-16 SNPs) in 7 individual FP outbreaks (**Fig. 3A**). These findings are consistent with the hypothesis that contaminated food is the main source of these *C. perfringens* food poisoning outbreaks, which included all meat-based food stuffs e.g. cooked sliced beef, lamb, chicken curry, cooked turkey and cooked meat (**Table S1**).

**Fig. 3.**
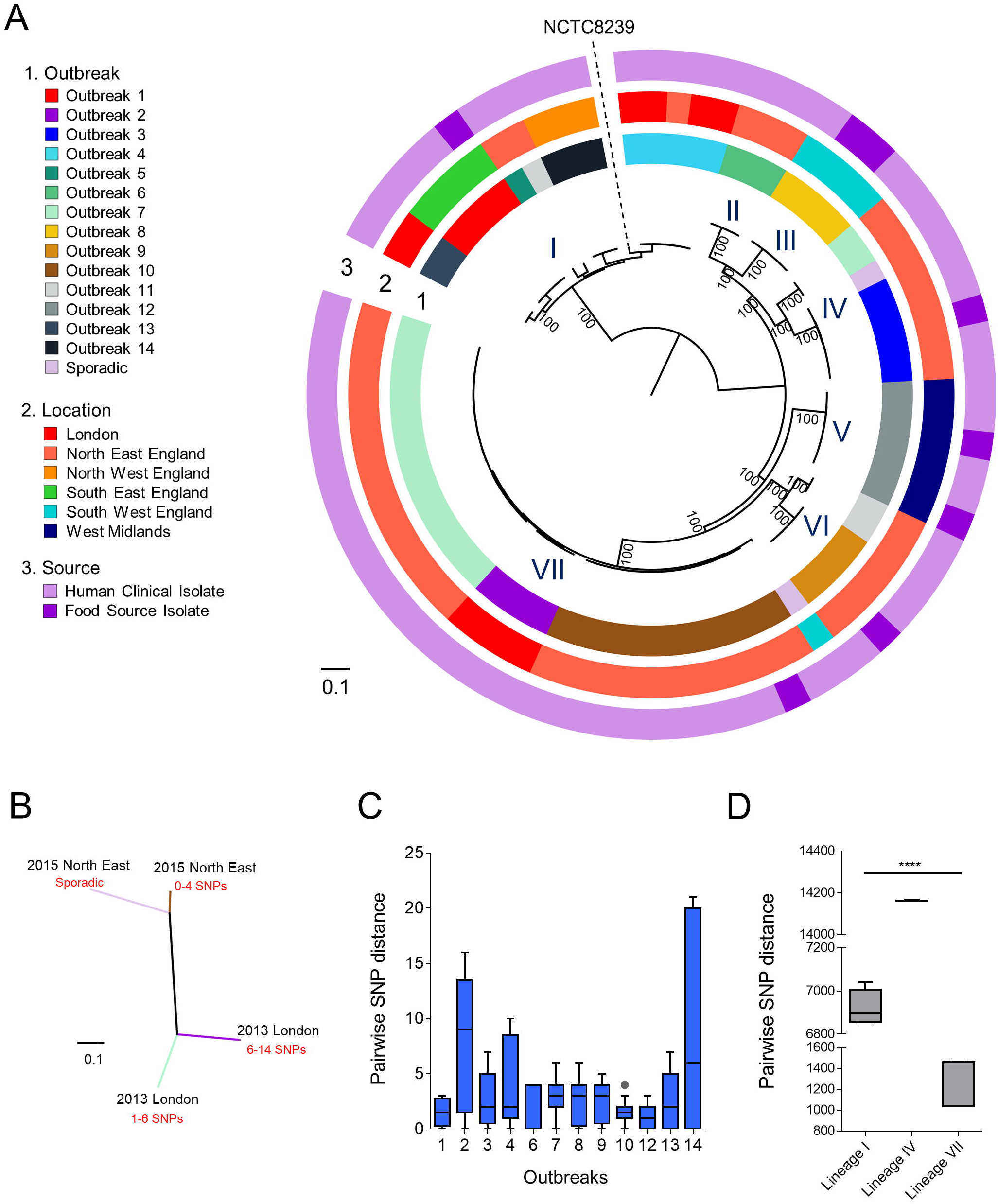
Phylogenomic analysis of 75 foodborne-associated *C. perfringens* isolates. **(A)** Mid-point rooted maximum-likelihood phylogeny of food-poisoning *C. perfringens* inferred from 70 613 SNPs (in core gene alignment) identified in 75 individual isolates. NCTC8239, a food poisoning strain isolated in 1949 (encodes *cpe* gene) is a RefSeq public reference genome. Lineages were determined via hierBAPS clustering analysis. Bootstrap values are represented in the tree. **(B)** Unrooted maximum-likelihood tree inferred from 2 505 SNPs in lineage VII (31 isolates). Four distinct clusters are identified in different outbreaks comprising genetically-similar strains (labeled in locations and years, and SNP range in outbreaks; branches are colour-coded corresponding to outbreak labels). **(C)** Pairwise core-SNP distance comparison in between isolates within outbreaks. **(D)** Pairwise core-SNP comparisons of within-major-lineage isolates in between individual outbreaks. Lineage I: outbreaks 1,4,13 and 14; Lineage IV: outbreaks 3 and 7; Lineage VII: outbreaks 2, 7 and 10. Data: Kruskal-Wallis test; **** P<0.0001.

### Virulence gene content

Diarrhoea symptoms associated with *C. perfringens* are primarily due to production of the pore-forming toxin enterotoxin (CPE) by *C. perfringens* type F strains (2, 60). Additional virulence determinants implicated in diarrhoea include sialidase (NanI), which is linked to enhanced intestinal attachment and an accessory role in enhancing CPE cell-toxicity, and also pore-forming toxin perfringolysin (PFO), a toxin known to act synergistically with alpha-toxin (phospholipase produced by all *C. perfringens* strains) to inflict intestinal cell damage (24, 61, 62). Moreover, antibiotic-resistant *C. perfringens* are reported to be prevalent, particularly within poultry, thus antimicrobial resistance (AMR) profiles of *C. perfringens* may be linked with prolonged *C. perfringens* associated-infections, and may hamper downstream treatments strategies (63, 64). To probe these important virulence-associated traits we screened isolates for toxin and AMR genes, based on both genome assemblies and raw sequence reads.

Enterotoxin gene *cpe* was detected in all, but 4 isolates (PH017, PH029, PH045 and PH156 were *cpe*-negative), which was confirmed by PCR, with the exception of PH029 which was initially determined to be *cpe*-positive via PCR (96.4%; **Table S1; Fig. S1B**).

CH isolates (average 9.6 ± 1.0 toxin genes per isolate) encoded significantly more toxin genes (P<0.001) than FP isolates (7.3 ± 1.9 toxin genes per isolate; **Fig. S1A**). CH isolates in lineages II-IV generally possessed identical toxin profiles (**Fig. 4A**); colonisation-related sialidase-encoding genes *nanI, nanJ* and *nagH*, haemolysin PFO gene *pfo*, and *cpb2* (**Fig. S1C-G**), which produces a vital accessory toxin beta-2 toxin (CPB2) associated with CPE-mediated pathogenesis (65). However, CH isolates did not harbour many acquired AMR genes; only 6 isolates (out of 35; 17%) encoded tetracycline resistant genes *tet(P)*, one isolate encoded aminoglycoside resistant gene *APH(3′)*, with remaining isolates not encoding any acquired AMR genes, other than the intrinsic AMR gene *mprF*.

**Fig. 4.**
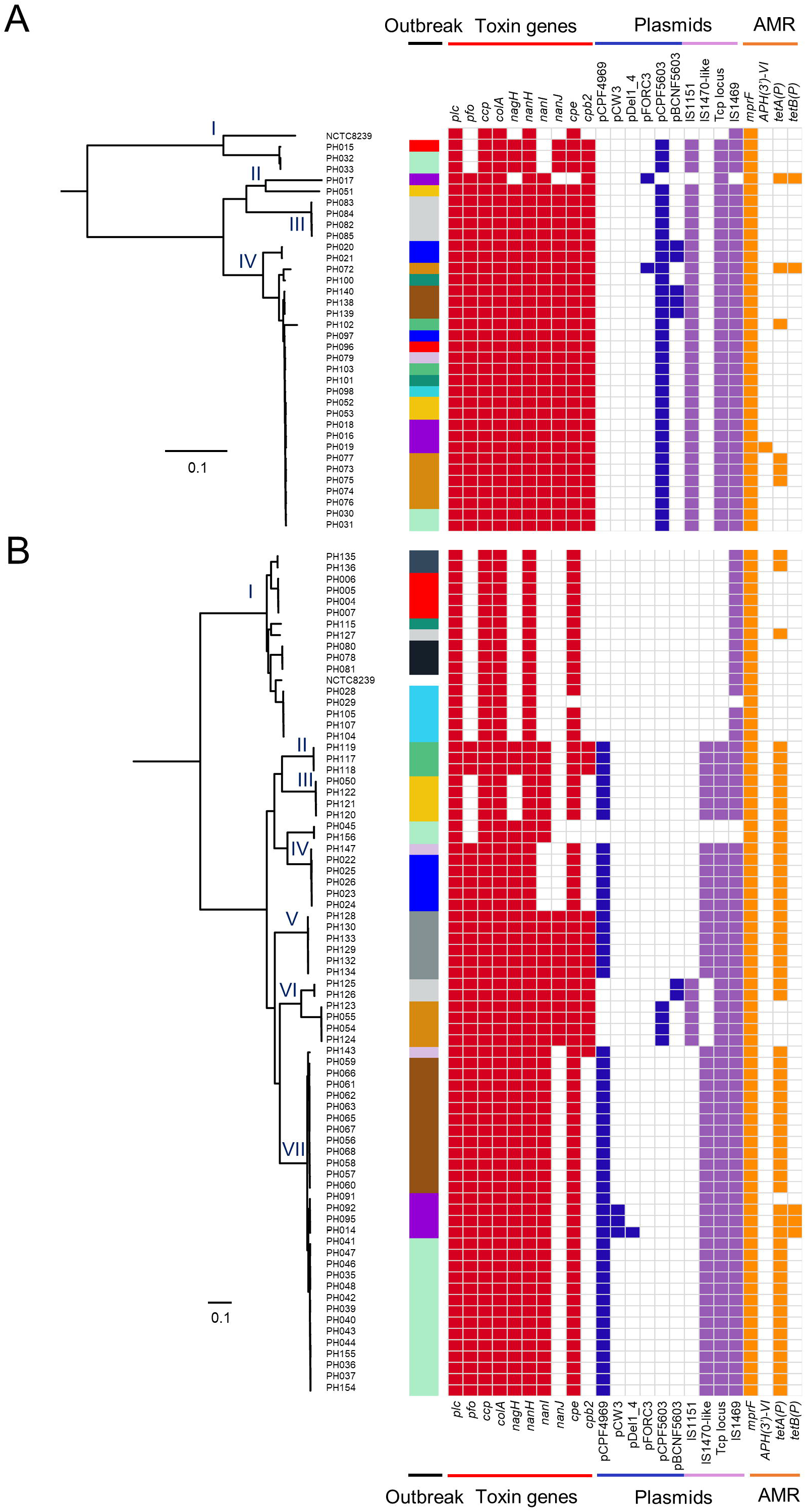
Full virulence profiles of *C. perfringens* isolates including virulence plasmids. Binary heatmaps displaying presence and absence of toxin genes, AMR genes, plasmids, plasmid-related sequences and *tcp* conjugative loci in corresponding isolates: **(A)** CH isolates **(B)** FP isolates. Outbreaks were colour-coded according to the colour system in previous figures. Coloured cells represent presence of genes and white cells represent absence of genes. Heatmaps were generated in R.

FP isolates had a more variable virulence gene profile (**Fig. 4B**). Isolates in lineage I had identical toxin genes, including *cpe*, but these isolates did not encode toxins such as PFO, CPB2 and sialidase NanI, and only three isolates in this lineage carried tetracycline resistant genes (19%). Most isolates within lineages II-VII (92%; 53/58) encoded *tetA(P)*, with this AMR gene significantly enriched in all FP isolates (74.3%; 55/74; P<0.0001; **Fig. S1H**), when compared to CH isolates (17.1%; 6/35). Furthermore, most isolates in FP lineages II-VII also encoded toxin genes *cpe, nanI* and *pfo*, and 16 isolates (28%) possessed the accessory toxin gene *cpb2*. Statistically these FP isolates (8.0 ± 1.5 toxin genes) encoded more toxin genes than those belonging to FP lineage I (4.9 ± 0.3 toxin genes; P<0.0001; **Fig. S1I**) which may suggest increased virulence. Initial plasmid detection indicated that virulence plasmid pCPF5603 was associated with CH isolates (97% in CH plasmid-carrying isolates; 34/35; P<0.0001), while plasmid pCPF4969 was linked to FP isolates (86.4% in FP plasmid-carrying isolates; 51/59; P<0.0001; **Fig. S1J**).

### Specific plasmid-associated lineages and potential CPE-plasmid transmission

The CPE toxin is responsible for the symptoms of diarrhoea in food poisoning, and non-foodborne illnesses, in the latter usually lasting >3 days, and up to several weeks (2, 66). Genetically, whilst chromosomal encoded *cpe* strains are primarily linked to food-poisoning (67, 68), non-foodborne diarrhoea is usually associated with plasmid-borne *cpe* strains (66, 69, 70). We performed an in-depth plasmid prediction on our datasets and analysis indicated that CH isolates predominantly harboured pCPF5603 plasmids (34/35 isolates; 97%) encoding *cpb2* and *cpe* genes, whilst FP isolates carried primarily pCPF4969 plasmids (45/75 isolates; 60%) encoding *cpe* but not *cpb2* genes (**Fig. S1J**). We also performed a genome-wide plasmid-specific sequence search to confirm our findings including IS *1151* (pCPF5603), IS *1470*-like (pCPF4969) and plasmid conjugative system *tcp*genes (**Fig. 4A-B**) (71–73).

To further examine and confirm the predicted plasmids, we extracted plasmid sequences (complete unassembled single contig) from three isolates per CH or FP group, and compared with reference plasmids (**Fig. 5A-C**). The extracted plasmid sequences closely resembled the respective reference plasmids, with near-identical nucleotide identity (>99.0%), plasmid size and GC content (**Fig. 5B-C; Table S9**), thus supporting the findings that these two intact plasmids (pCPF4969 and pCPF5603) are present in these isolates.

**Fig. 5.**
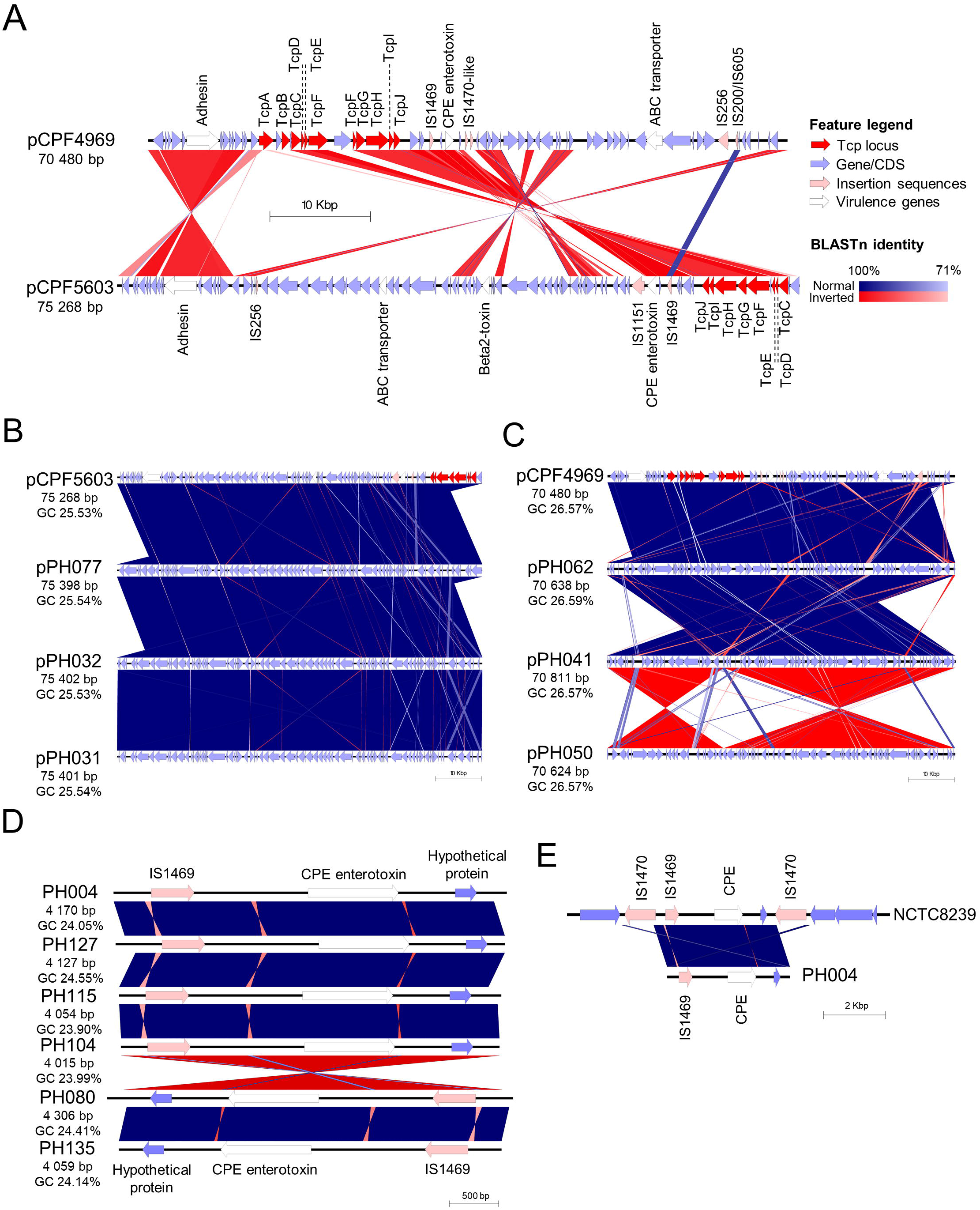
Investigations on predicted plasmids carried by CH and FP isolates. **(A)** Comparative genomic visualisation of reference plasmids pCPF4969 and pCPF5603 with annotated features. **(B)** Genomic comparison of pCPF5603 reference plasmid and predicted plasmids from CH genomes. **(C)** Plasmid comparison in between pCPF4969 plasmid and FP-isolate predicted plasmids. **(D)** CPE-regions (Tn*5565*) extracted computationally from FP lineage I representative isolate genomes. **(E)** A computationally extracted 11-kbp region of NCTC8239 that encodes Tn5565 (including cpe and flanking IS*1470* elements) compared to predicted Tn*5565* from PH004. Figures were produced using Easyfig v2.2.2.

Although chromosomal-*cpe* strains are considered as the primary strain type to be associated with food poisoning, our dataset demonstrated that plasmid-*cpe C. perfringens* strains were predominantly associated with food poisoning (82.6%; 90/109), with only 17.4% FP isolates encoding a copy of *cpe* on the chromosome (no plasmid detected). Putatively, plasmid transfer may have occurred in CH outbreak 7 isolates (n=4; PH030, PH031, PH032, PH033), as two isolates reside within lineage IV, whilst the other two isolates nest within the genetically distant CH lineage I (genetic distance >10 000 SNPs), however all 4 isolates harboured plasmid pCPF5603 (**Fig. 4A**). CH outbreaks 1 and 8 also had dissimilar strains (nested within separate lineages) with identical plasmids. This analysis denotes that multiple distinct strains, but carrying the same *cpe*-plasmid, may be implicated in these CH outbreaks, with previous work showing *in vitro* plasmid transfer among *C. perfringens* strains via conjugation (*cpe*-positive to *cpe*-negative strains) (74).

Previous studies have demonstrated that *C. perfringens* with chromosomally-encoded *cpe* are genetically divergent from plasmid-*cpe* carriers. Within the FP phylogeny there was a distinct lineage of isolates (lineage I; n=17) that appear to encode *cpe* chromosomally. These isolates had significantly smaller genomes (genome size 2.95 ± 0.03 Mb vs 3.39 ± 0.08 Mb outside-lineage; n=93; P<0.0001; **Table S10**), were most similar to reference genome NCTC8239 (ANI≥99.40%) and appeared to lack plasmids. This was further evidenced by historical chromosomal-*cpe* strain NCTC8239 nesting in this lineage with these newly sequenced strains (FP lineage I; **Fig. 4**) (70, 72, 75). To further investigate this hypothesis, the *cpe*-encoding region (complete single contig from high-coverage assemblies) was extracted from representative isolates in lineage I (n=6), and comparative genomics was performed (**Fig. 5D**). These consistently smaller (~4.0-4.3 kb) contigs were almost identical in nucleotide identity (>99.9%) when compared with the *cpe*-encoding region of chromosomal-*cpe* strain NCTC8239, confirming that these isolates possessed the same *cpe* genomic architecture as NCTC8239 and confirmed as transposable element Tn*5565* (**Fig. 5E**). In addition, PH029 was the only outbreak isolate not detected to encode *cpe* within the lineage I outbreak cluster, despite having a clonal relationship with PH028, PH104, PH105 and PH107 (FP outbreak 4; **Fig. 4B**). This suggests Tn*5565* loss may have occurred due to extensive sub-culturing (this is supported by initial PCR results being *cpe*-positive; see **Table S1**). Analysis also indicates that *cpe* was closely associated with IS *1469* independent of where it was encoded, as this insertion sequence was detected exclusively in all *cpe*-encoding genomes (100%; **Fig. 4A-B**).

### Accessory genome virulence potentials

The 110 strain *C. perfringens* pangenome consisted of 6 219 genes (including NCTC8239; **Fig. S2**); 1 965 core genes (31.5%), and 4 254 accessory genes (68.5%); with ~30-40% genes in any individual strain encoded within the accessory genome, potentially driving evolution and genome restructuring. Mobile genetic elements including plasmids, genomic islands and prophages could potentially contribute to virulence, given the plasticity of the genome. To explore this in more detail, we further analysed the accessory genomes, comparing different sub-sets of *C. perfringens* isolates. We first identified subset-specific genes using a bacterial pan-GWAS approach, with these genes further annotated based on NCBI RefSeq gene annotations and categorised under COG classes into three comparison groups: (1) CH vs FP; (2) FP outbreaks; (3) FP lineage I, FP lineage II-VII and CH-FP plasmid-CPE isolates (**Fig. S3A-C**).

Phosphotransferase system (PTS)-related genes (n=4) were encoded exclusively in CH isolates (present in 26/35 CH isolates; **Fig. S3A and Table S11**). These genes may contribute to the isolates’ fitness to utilise complex carbohydrates (COG category G) in competitive niches, like the gastrointestinal tract (76). PTS genes have been linked to virulence regulation in other pathogens including foodborne pathogen *Listeria monocytogenes* (77). Heat-shock protein (Hsp70) DnaK co-chaperone was annotated in FP-specific accessory genome (present in 57/74 FP isolates), which may be involved in capsule and pili formation which may facilitate host colonisation (78–80).

Accessory genes specific to each FP outbreaks were variable (**Fig. S3B-C and Table S12**), but 3 annotated functional classes were conserved; L (replication, recombination & repair), M (cell wall/membrane/envelope biogenesis) and V (defense mechanisms). Prominent genes detected in all isolates included phage-related proteins (n=49) (L, M and S), glycosyltransferases (n=37) (M), restriction modification systems (n=16) (V), transposases (n=9) (L) and integrases (n=8) (L). It was evident that most genes were associated with phages, seemingly a major source of mobile genetic transfer.

Correspondingly, less group-specific accessory genes where present compared with other isolates in lineages II-VII (**Fig. S3C and Table S13**). Notably, multidrug transporter ‘small multidrug resistance’ genes were exclusively detected in FP lineage I isolates, whereas ABC transporters were more commonly encoded in plasmid-carrying isolates (virulence plasmids pCPF5603 and pCPF4969 carry various ABC transporter genes). The Mate efflux family protein gene was detected solely in lineage II-VII isolates.

### Prophage genomes linked to enhanced *C. perfringens* fitness

Phage are important drivers of bacterial evolution and adaptation, and presence of prophage within bacterial genomes is often associated with enhanced survival and virulence e.g. sporulation capacity and toxin secretion (81–83). Thus, mining phages in foodborne *C. perfringens* genomes could reveal insights into the role of bacteriophage in modulating diversity and pathogenesis traits (25). We identified through PHASTER a total of 7 prophages in all 109 genomes (**Fig. S4A-B**). Further exploration into virulence and survival-enhancing genes (**Fig. S4C**) encoded in these predicted prophage regions revealed the presence of virulence-related enzyme sialidase NanH (promotes colonisation), putative enterotoxin EntB, various ABC transporters (linked to multidrug resistance) and toxin-linked phage lysis holin (probable link to toxin secretion)(61, 84–86). No differences in number of prophages carried were detected between CH and FP isolates (**Fig. S4D-E**). These data suggested that phages could potentially contribute to increased accessory virulence within the genomes of food-poisoning associated *C. perfringens*, and indicates further research, using both experimental and genomic approaches, is required.

## Discussion

*C. perfringens* is often associated with self-limiting or longer-term gastroenteritis, however our knowledge on the genomic components that may link to disease symptoms or epidemiological comparisons between outbreaks is limited. In this study, WGS data and in-depth genomic analysis on a representative sub-set of 109 gastrointestinal outbreak-associated *C. perfringens* isolates, revealed potential epidemic phylogenetic clusters linked to plasmid carriage, and specific virulence determinants, which were strongly associated with outbreak isolates.

In the context of disease control it is important to gain detailed genomic information to predict transmission modes for pathogens. Our analysis of care home isolates indicated a specific persistent clone may have been responsible for up to 9 individual gastrointestinal outbreaks in North East England over the 2013-2017 period, which represents the majority reported gastroenteritis outbreaks (>80%) in this area (10). Interestingly, a previous study indicated presence of persistent identical *C. perfringens* genotypes within care home settings, with several individuals harbouring identical strains (as determined via PFGE profiling) throughout a 9-month sampling period, however none of these isolates were positive for the *cpe* gene (87). Furthermore, although care home isolates were defined as ‘non-foodborne’ according to local epidemiological investigations as no food samples were identified as *C. perfringens*-positive, these outbreaks may have resulted from contaminated food products not sampled. Indeed, a recent investigation into fresh meat products (>200 samples) demonstrated significant contamination; beef (~30%), poultry and pork (both ~26%), with 90% of strains *cpe*-positive, suggesting food chain(s) or farmed animals as potential reservoirs of enterotoxigenic *C. perfringens* (70, 88). Interestingly, 18% prevalence of *cpe*-positive *C. perfringens* strains had previously been reported in food handlers’ faeces (confirmed via PCR), denoting a potential role of the human reservoir in outbreaks (89). Determination of these potential reservoirs in the spread of *cpe*-positive *C. perfringens* to at-risk populations would necessitate a One Health approach, and large-scale WGS-based screening to ascertain the phylogenomics of strains isolated from diverse sources surrounding outbreak-linked vicinities.

Successful colonisation of invading *C. perfringens* is required for efficient toxin production, which ultimately leads to gastrointestinal symptoms. Through computational analysis we determined that plasmid-*cpe* (specifically plasmids pCPF4969 and pCPF5603) carrying strains predominated within both non-foodborne and food-poisoning outbreak-related *C. perfringens* isolates (~82%). These two virulence plasmids, pCPF4969 and pCPF5603, encoded several important virulence genes including ABC transporter and adhesin (also known as collagen adhesion gene *cna*) that could contribute to enhanced survival and colonisation potential of *C. perfringens* within the gastrointestinal tract (90). Plasmid pCPF4969 also contained a putative bacteriocin gene that may allow *C. perfringens* to outcompete other resident microbiota members, and thus overgrow and cause disease in the gut environment (73). Plasmid pCPF5603 encoded important toxin genes *cpe* and *cpb2*, plus additional toxins, many of which are linked to food poisoning symptoms, such as diarrhoea and cramping.

Interestingly, 4 out of 109 outbreak-associated strains were *cpe*-negative, suggesting secondary virulence genes (e.g. *pfo* and *cpb2*) may be associated with *C. perfringens-associated* gastroenteritis. A recent WGS-based study on FP *C. perfringens* outbreaks in France determined that ~30% of isolates were *cpe*-negative (13/42) (26), indicating this gene may not be the sole virulence determinant linked to *C. perfringens* gastroenteritis. Although we observed less *cpe*-negative strains in our collection in comparison to this study, this may be due to our targeted *cpe*-positive isolation strategy (standard practice at PHE). Thus, to determine the importance and diversity of *cpe*-negative strains in FP outbreaks this will require untargeted isolation schemes in the future.

Typical *C. perfringens-associated* food poisoning was previously thought to be primarily caused by chromosomal-*cpe* strains. This is linked to their phenotypic capacity to withstand high temperatures (via production of a protective small acid soluble protein), and high salt concentrations during the cooking process, in addition to the shorter generation time, when compared to plasmid-*cpe* carrying strains (68, 91). Previous studies have indicated that these strains commonly assemble into distinct clusters that lack the *pfo* gene, which we also noted in the FP lineage I data from this study (26, 92–94). Nevertheless, plasmid-borne *cpe*-carrying strains (pCPF4969 or pCPF5603) have also been associated with previous food poisoning outbreaks, with a previous study indicating that pCPF5603-carrying strains (encoding IS *1151*) were associated with food poisoning in Japanese nursing homes (7 out of 9 isolates) (71). However, these plasmid-*cpe* outbreaks have been described as a relatively uncommon occurrence, thus it is surprising that our findings indicate that most outbreak isolated strains (81.6%; 89/109) carried a *cpe*-plasmid (60, 67). The fact that plasmid-*cpe* strains can cause diverse symptoms including short-lived food poisoning, and long-lasting non-foodborne diarrhoea, implicates additional factors in disease pathogenesis. The gut microbiome may be one such host factor as previous studies have reported that care home residents have a less diverse and robust microbiota when compared to those residing in their own homes (including individuals colonised with *C. perfringens)*, and thus impaired ‘colonisation resistance’ may mean certain *C. perfringens* strains can overcome these anti-infection mechanisms and initiate disease pathogenesis (64, 87, 95).

Chromosomal-*cpe* is reported to be encoded on a transposon-like element Tn*5565* (6.3 kb, with flanking copies of IS *1470*), which can form an independent and stable circular-form in culture extracts (losing both copies of IS *1470*) (72, 74). This transposition element TN*5565* was commonly thought to be integrated into the chromosome at a specific site as a unit. The fact that our computational analysis failed to detect any *cpe*, IS *1469* (*cpe*-specific), and IS *1470* (Tn*5565*-specific) in the high-sequencing-coverage PH029 genome (317X sequencing depth/coverage) indicates that Tn*5565* can be lost or may be passed on to other *C. perfringens* cells. However, it should be noted that flanking IS *1470* of Tn*5565* may not have been correctly assembled during the genome assembly process due to the repetitive nature of those sequences (short-read sequencing).

As WGS provides enhanced resolution to identify outbreak-specific clonal strains, our study highlighted the importance of implementing WGS for *C. perfringens* profiling in reference laboratories, in place of the conventional fAFLP (92, 93, 96). Routine *C. perfringens* surveillance of the care home environment and staff could prove critical for vulnerable populations, as outbreaks could rapidly spread, and this approach could potentially pinpoint the sources of contamination, and eventually eliminate persistent *cpe*-strains in the environment (87). In light of the potential rapid transmissibility of *C. perfringens cpe*-strains responsible for food-poisoning outbreaks, real-time portable sequencing approaches such as the MinION, could facilitate the rapid identification of outbreak strains, which has been recently been reported to identify outbreak *Salmonella* strains in <2h (97, 98).

Our data highlights the genotypic and epidemiology relatedness of a large collection of *C. perfringens* strains isolated from food poisoning cases from across England and Wales, and indicates potential circulation of disease-associated strains, and the potential impact of plasmid-associated-*cpe* dissemination, linked to outbreak cases. This study indicates that further WGS phylogenetic and surveillance studies of diversely-sourced *C. perfringens* isolates are required for us to fully understand the potential reservoir of food poisoning-associated strains, so that intervention or prevention measures can be devised to prevent the spread of epidemiologically important genotypes, particularly in vulnerable communities, including older adults residing in care homes.

## Electronic supplemental materials

Supplementary Figure S1

Supplementary Figure S2

Supplementary Figure S3

Supplementary Figure S4

Supplementary Table S1

Supplementary Table S2

Supplementary Table S3

Supplementary Table S4

Supplementary Table S5

Supplementary Table S6

Supplementary Table S7

Supplementary Table S8

Supplementary Table S9

Supplementary Table S10

Supplementary Table S11

Supplementary Table S12

Supplementary Table S13

## Acknowledgments

This work was supported by a Wellcome Trust Investigator Award (100974/C/13/Z), and the Biotechnology and Biological Sciences Research Council (BBSRC); Institute Strategic Programme Gut Microbes and Health BB/R012490/1, and its constituent project(s) BBS/E/F/000PR10353 and BBS/E/F/000PR10356, and Institute Strategic Programme Gut Health and Food Safety BB/J004529/1 to LJH, and Institute Strategic Programme Microbes in the Food Chain BB/R012504/1 to AEM. This research was supported in part by the NBI Computing infrastructure for Science (CiS) group through the provision of a High-Performance Computing (HPC) Cluster. We also thank Dr. Andrew Page (Quadram Institute, UK) for the helpful discussion on computational analysis.

R.K., C.A. and L.J.H. designed the study. R.K. and A.P. processed the sequencing data. R.K. performed the genomic analysis and graphed the figures. S.C. and A. P. provided essential assistance in genome assembly and genomic analysis. A.M. contributed in genomic analysis tools and edited the manuscript, R.K., C.A. and L.J.H. analysed the data and co-wrote the manuscript. CS and CA managed the culture collection, genotyping, clinical data collection, and processed samples for sequencing.

## Footnotes

1 https://www.sanger.ac.uk/resources/downloads/bacteria/nctc/

