## Supplementary figures and images for "Phylogenomic analysis of *Clostridium perfringens* identifies isogenic strains in gastroenteritis outbreaks, and novel virulence-related features"

### Supplementary Figure 1

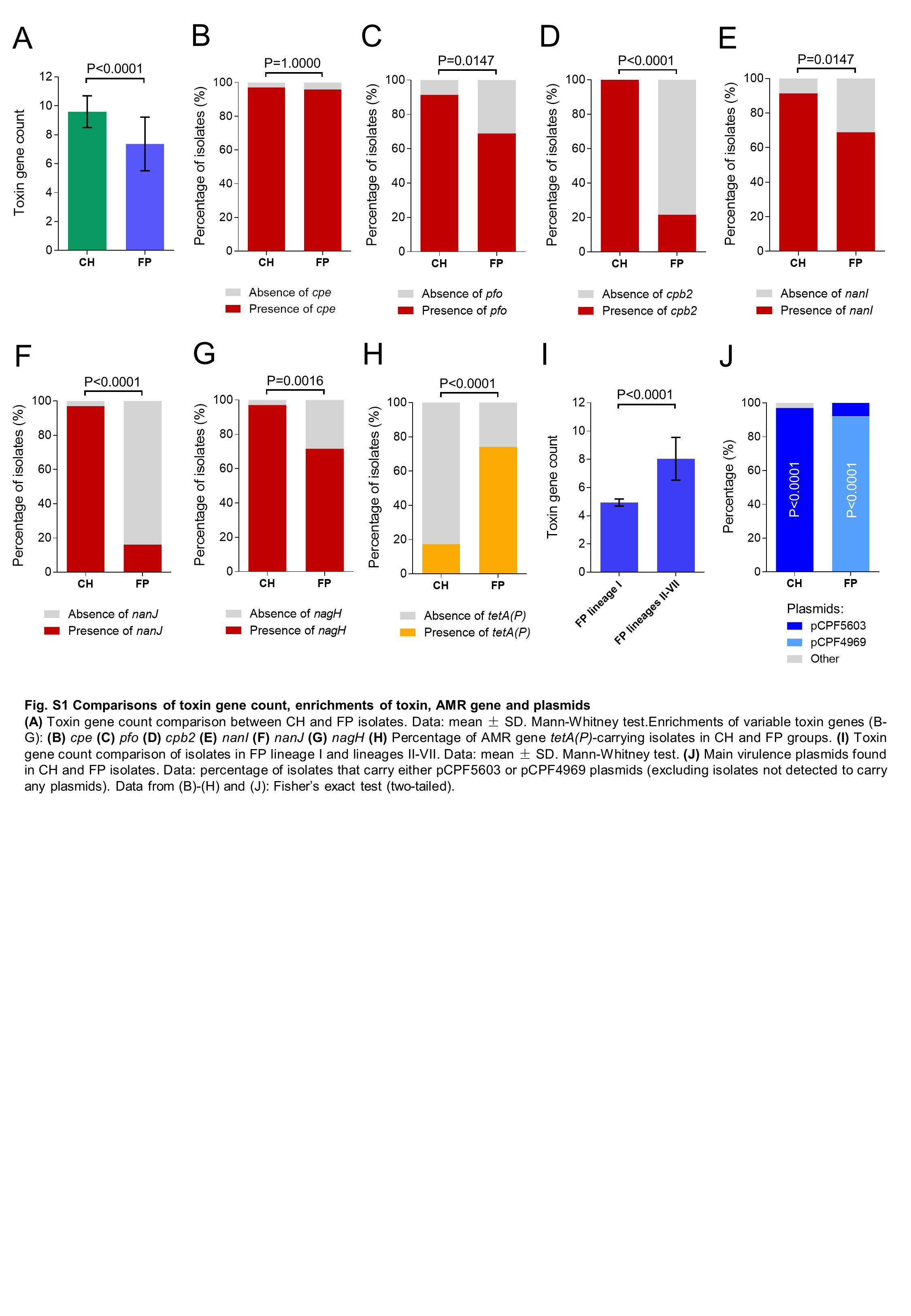

### Supplementary Figure 2

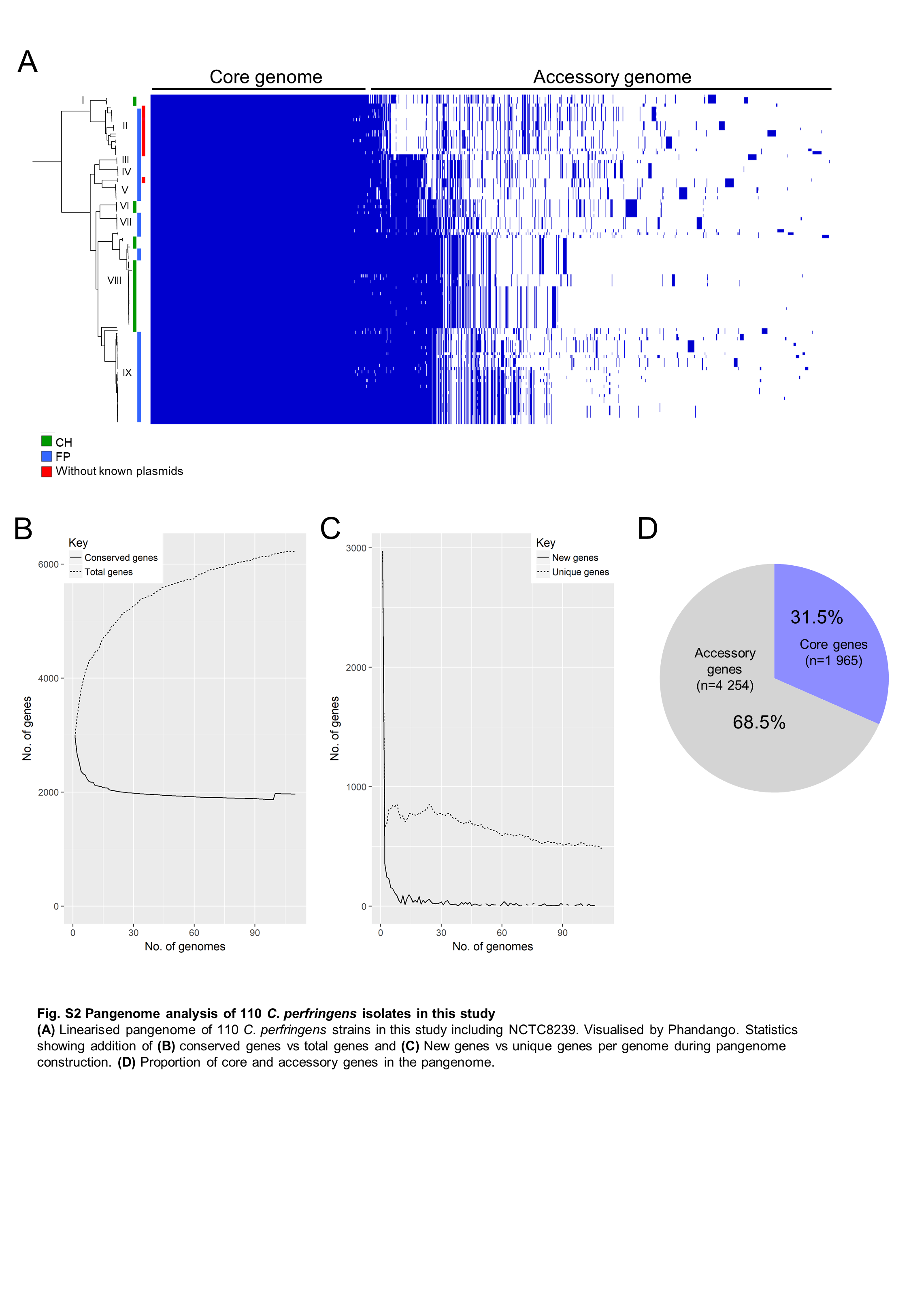

### Supplementary Figure 3

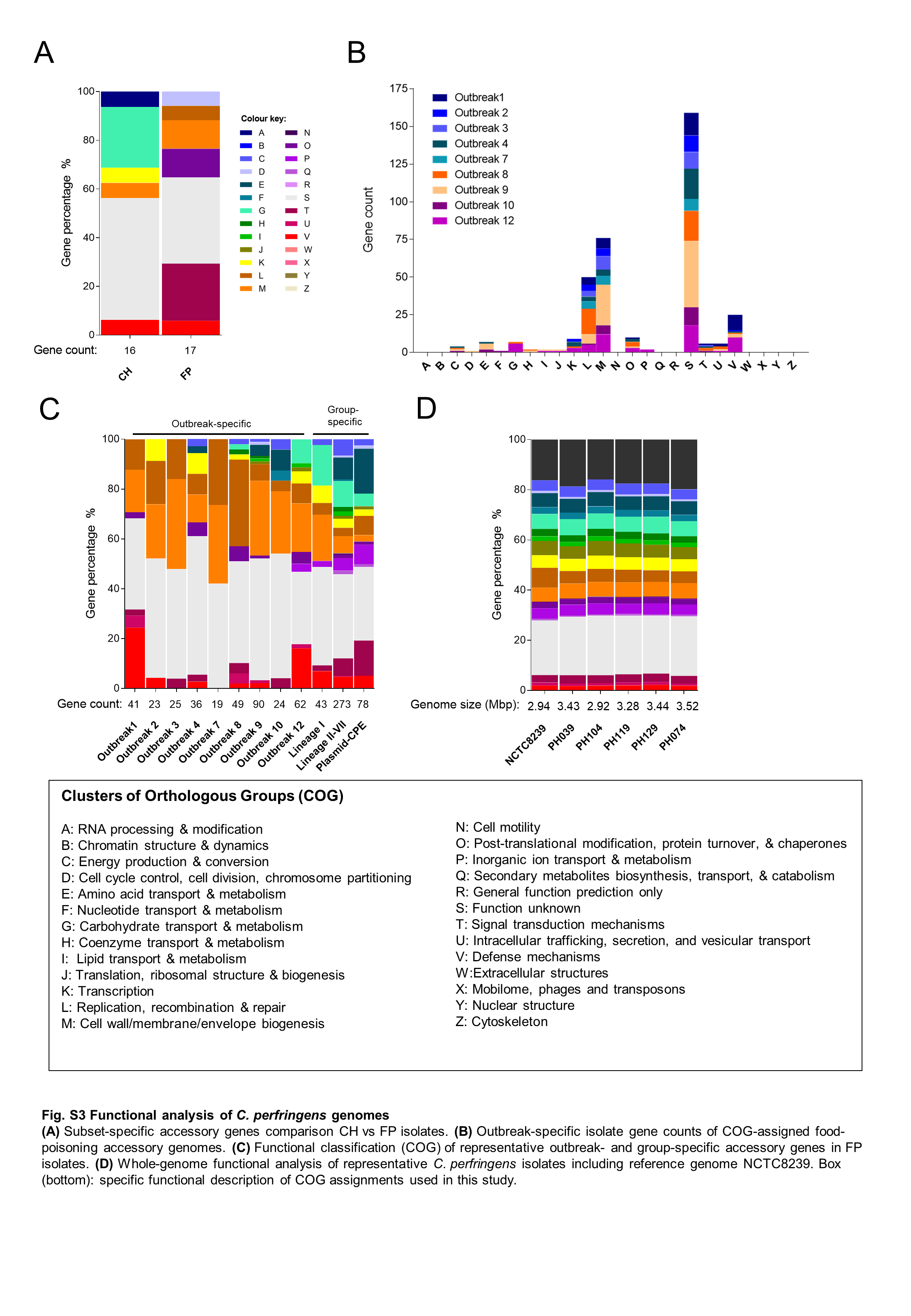

### Supplementary Figure 4

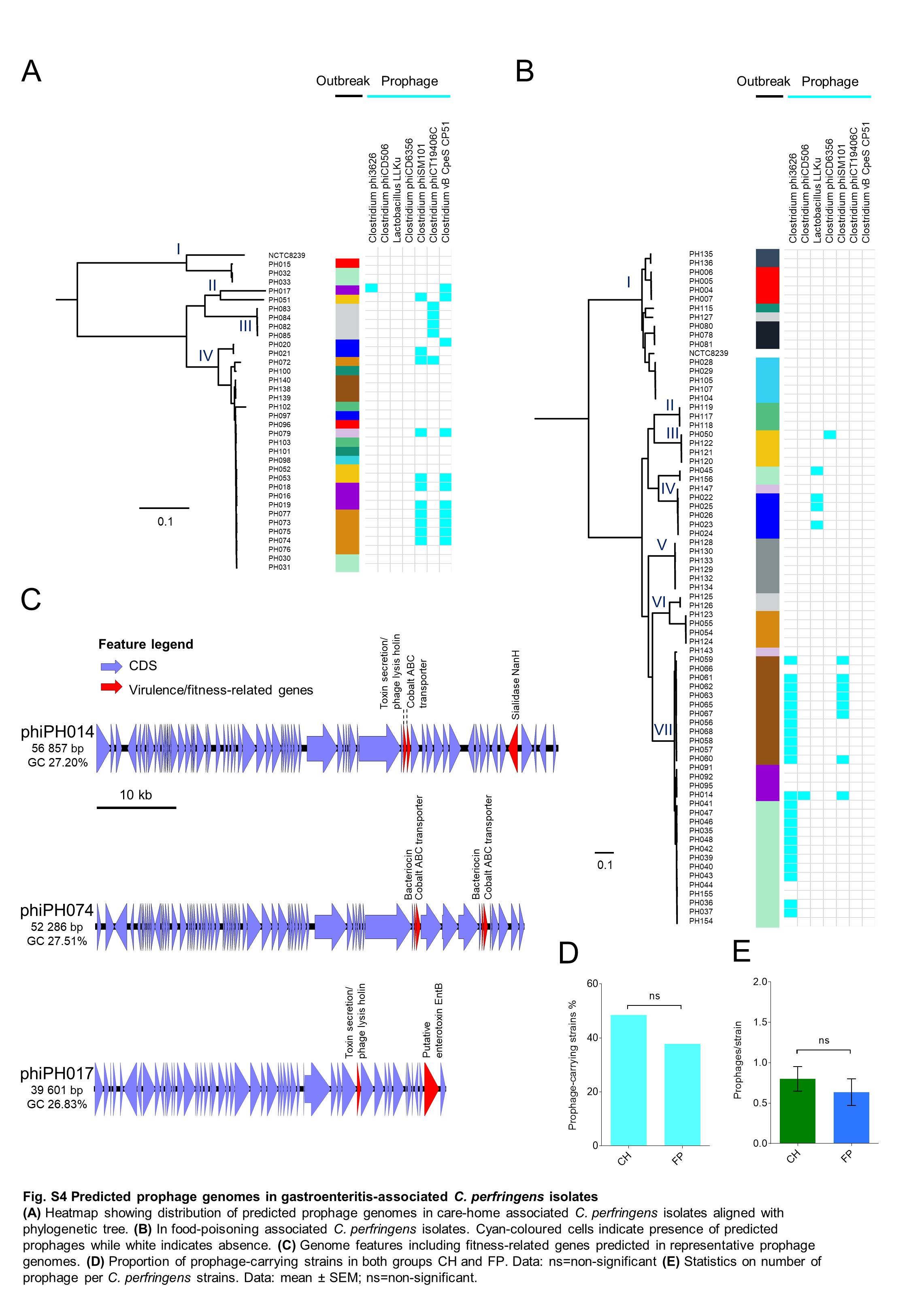
