## Supplementary Table 1 for "Phylogenomic analysis of *Clostridium perfringens* identifies isogenic strains in gastroenteritis outbreaks, and novel virulence-related features"

**Table S1.** Clinical and genomic (assembly and annotation) data of CH-associated (CH) *C. perfringens* isolates in this study. ANI (Average Nucleotide Identity) was compared with reference genome (RefSeq) *C. perfringens* NCTC8239. Cohort: CH represents care home-associated; FP denotes general food poisoning isolates.

|  | **Clinical metadata** | | | | | | **Sequencing information** | | | | **Genome assembly and annotation statistics** | | | | | | | | **PCR** |
| --- | --- | --- | --- | --- | --- | --- | --- | --- | --- | --- | --- | --- | --- | --- | --- | --- | --- | --- | --- |
|  | **Isolate** | **Source** | **Year** | **Region** | **Cohort** | **Out-break** | **Illumina sequencing platform** | **Read length (bp)** | **Reads (bp)** | **Sequencing coverage (X)** | **GC (%)** | **Contig number** | **Genome Size (bp)** | **tRNA** | **rRNA** | **CDS** | **Gene** | **ANI** | **PCR *cpe*** |
| 1 | PH015 | Individual | 2013 | North east | CH | 1 | HiSeq | 151 | 3854793 | 377 | 28.01 | 100 | 3081476 | 84 | 13 | 2731 | 2829 | 98.70 | positive |
| 2 | PH096 | Individual 1 | 2013 | North east | CH | 1 | MiSeq | 101 | 1267411 | 74 | 28.08 | 239 | 3422714 | 58 | 7 | 3105 | 3171 | 97.32 | positive |
| 3 | PH016 | Individual 1 | 2014 | North east | CH | 2 | HiSeq | 151 | 2853518 | 244 | 28.07 | 66 | 3518998 | 84 | 12 | 3206 | 3303 | 97.29 | positive |
| 4 | PH017 | Individual 2 | 2014 | North east | CH | 2 | HiSeq | 151 | 2625370 | 240 | 28.18 | 23 | 3291535 | 72 | 13 | 2935 | 3021 | 97.15 | negative |
| 5 | PH018 | Individual 2 | 2014 | North east | CH | 2 | HiSeq | 151 | 2325775 | 202 | 28.1 | 26 | 3475247 | 79 | 11 | 3164 | 3255 | 97.30 | positive |
| 6 | PH019 | Individual 3 | 2014 | North east | CH | 2 | HiSeq | 151 | 2615637 | 226 | 28.14 | 25 | 3487876 | 87 | 11 | 3170 | 3269 | 97.30 | positive |
| 7 | PH020 | Individual 1 | 2014 | North east | CH | 3 | HiSeq | 151 | 2483432 | 220 | 28.05 | 33 | 3397222 | 75 | 11 | 3075 | 3162 | 97.29 | positive |
| 8 | PH021 | Individual 2 | 2014 | North east | CH | 3 | HiSeq | 151 | 3350806 | 293 | 28 | 31 | 3445998 | 74 | 11 | 3144 | 3230 | 97.28 | positive |
| 9 | PH097 | Individual 3 | 2014 | North east | CH | 3 | MiSeq | 101 | 1138038 | 68 | 28.12 | 398 | 3371290 | 29 | 3 | 3049 | 3082 | 97.36 | positive |
| 10 | PH098 | Individual 1 | 2014 | North east | CH | 4 | MiSeq | 101 | 1500399 | 87 | 28.08 | 245 | 3454553 | 58 | 6 | 3137 | 3202 | 97.34 | positive |
| 11 | PH100 | Individual 1 | 2014 | North east | CH | 5 | MiSeq | 101 | 1457550 | 87 | 28.15 | 184 | 3354426 | 56 | 8 | 3010 | 3075 | 97.32 | positive |
| 12 | PH101 | Individual 2 | 2014 | North east | CH | 5 | MiSeq | 101 | 2004838 | 116 | 28.04 | 87 | 3481641 | 60 | 7 | 3179 | 3247 | 97.30 | positive |
| 13 | PH102 | Individual 1 | 2014 | North east | CH | 6 | MiSeq | 101 | 1476804 | 87 | 28.04 | 311 | 3420587 | 60 | 7 | 3110 | 3178 | 97.32 | positive |
| 14 | PH103 | Individual 2 | 2014 | North east | CH | 6 | MiSeq | 101 | 860585 | 50 | 28.07 | 274 | 3448541 | 42 | 5 | 3128 | 3176 | 97.33 | positive |
| 15 | PH030 | Individual 1 | 2014 | North east | CH | 7 | HiSeq | 151 | 2567442 | 226 | 28.07 | 27 | 3427777 | 67 | 8 | 3092 | 3168 | 97.30 | positive |
| 16 | PH031 | Individual 1 | 2014 | North east | CH | 7 | HiSeq | 151 | 2606506 | 229 | 28.08 | 21 | 3430445 | 74 | 10 | 3093 | 3178 | 97.31 | positive |
| 17 | PH032 | Individual 2 | 2014 | North east | CH | 7 | HiSeq | 151 | 2889707 | 277 | 27.94 | 109 | 3139693 | 78 | 13 | 2827 | 2919 | 98.70 | positive |
| 18 | PH033 | Individual 2 | 2014 | North east | CH | 7 | HiSeq | 151 | 2287790 | 218 | 27.99 | 105 | 3163229 | 70 | 9 | 2850 | 2930 | 98.70 | positive |
| 19 | PH051 | Individual 1 | 2015 | North east | CH | 8 | HiSeq | 151 | 2218172 | 192 | 28.09 | 27 | 3480399 | 92 | 11 | 3155 | 3259 | 97.21 | positive |
| 20 | PH052 | Individual 2 | 2015 | North east | CH | 8 | HiSeq | 151 | 2665002 | 231 | 28.07 | 34 | 3481680 | 74 | 8 | 3188 | 3271 | 97.30 | positive |
| 21 | PH053 | Individual 3 | 2015 | North east | CH | 8 | HiSeq | 151 | 2587606 | 224 | 28.09 | 25 | 3484752 | 95 | 8 | 3188 | 3292 | 97.31 | positive |
| 22 | PH072 | Individual 2 | 2016 | North east | CH | 9 | HiSeq | 151 | 2510560 | 217 | 28.01 | 59 | 3492601 | 86 | 12 | 3153 | 3252 | 97.27 | positive |
| 23 | PH073 | Individual 3 | 2016 | North east | CH | 9 | HiSeq | 151 | 2426321 | 207 | 28.07 | 35 | 3537414 | 92 | 10 | 3246 | 3349 | 97.30 | positive |
| 24 | PH074 | Individual 4 | 2016 | North east | CH | 9 | HiSeq | 151 | 2486225 | 213 | 28.07 | 26 | 3522802 | 93 | 11 | 3229 | 3334 | 97.31 | positive |
| 25 | PH075 | Individual 5 | 2016 | North east | CH | 9 | HiSeq | 151 | 2353413 | 200 | 28.08 | 29 | 3538037 | 95 | 13 | 3250 | 3359 | 97.30 | positive |
| 26 | PH076 | Individual 6 | 2016 | North east | CH | 9 | HiSeq | 151 | 1995409 | 174 | 28.05 | 23 | 3450640 | 83 | 11 | 3121 | 3216 | 97.31 | positive |
| 27 | PH077 | Individual 7 | 2016 | North east | CH | 9 | HiSeq | 151 | 2455240 | 212 | 28.11 | 23 | 3491708 | 89 | 10 | 3178 | 3278 | 97.30 | positive |
| 28 | PH138 | Individual 1 | 2016 | Wales | CH | 10 | MiSeq | 101 | 1559472 | 92 | 28.06 | 222 | 3401096 | 58 | 9 | 3056 | 3124 | 97.33 | positive |
| 29 | PH139 | Individual 2 | 2016 | Wales | CH | 10 | MiSeq | 101 | 1111879 | 66 | 28.04 | 97 | 3402225 | 60 | 11 | 3069 | 3141 | 97.30 | positive |
| 30 | PH140 | Individual 3 | 2016 | Wales | CH | 10 | MiSeq | 101 | 1264793 | 75 | 28.06 | 257 | 3386184 | 34 | 5 | 3043 | 3083 | 97.33 | positive |
| 31 | PH082 | Individual 1 | 2017 | North east | CH | 11 | HiSeq | 151 | 2961546 | 263 | 28.11 | 30 | 3394345 | 90 | 10 | 3065 | 3166 | 97.18 | positive |
| 32 | PH083 | Individual 2 | 2017 | North east | CH | 11 | HiSeq | 151 | 1865948 | 165 | 28.07 | 21 | 3410579 | 89 | 12 | 3088 | 3190 | 97.18 | positive |
| 33 | PH084 | Individual 3 | 2017 | North east | CH | 11 | HiSeq | 151 | 1964261 | 174 | 28.1 | 22 | 3392793 | 95 | 11 | 3067 | 3174 | 97.18 | positive |
| 34 | PH085 | Individual 4 | 2017 | North east | CH | 11 | HiSeq | 151 | 2764341 | 245 | 28.11 | 23 | 3394191 | 88 | 11 | 3064 | 3164 | 97.18 | positive |
| 35 | PH079 | Individual | 2017 | North east | CH | Sporadic | HiSeq | 151 | 2568032 | 221 | 28.1 | 26 | 3496302 | 91 | 10 | 3184 | 3286 | 97.3 | positive |
| 36 | PH007 | Curry goat | 2011 | South East | FP | 1 | HiSeq | 151 | 2776491 | 284 | 27.93 | 81 | 2946980 | 74 | 13 | 2660 | 2748 | 99.42 | positive |
| 37 | PH004 | Individual 1 | 2011 | South East | FP | 1 | HiSeq | 151 | 2522471 | 260 | 27.89 | 95 | 2923443 | 89 | 12 | 2633 | 2735 | 99.43 | positive |
| 38 | PH006 | Individual 3 | 2011 | South East | FP | 1 | HiSeq | 151 | 2763075 | 284 | 27.87 | 95 | 2928275 | 91 | 13 | 2635 | 2740 | 99.42 | positive |
| 39 | PH005 | Individual 2 | 2011 | South East | FP | 1 | HiSeq | 151 | 2827331 | 291 | 27.89 | 96 | 2934006 | 87 | 14 | 2640 | 2742 | 99.42 | positive |
| 40 | PH014 | Individual 2 | 2013 | London | FP | 2 | HiSeq | 151 | 2781207 | 245 | 28.11 | 42 | 3423911 | 89 | 11 | 3074 | 3175 | 97.19 | positive |
| 41 | PH092 | Individual 2 | 2013 | London | FP | 2 | MiSeq | 101 | 702080 | 41 | 28.12 | 378 | 3378829 | 38 | 6 | 3006 | 3051 | 97.22 | positive |
| 42 | PH095 | Individual 5 | 2013 | London | FP | 2 | MiSeq | 101 | 1024205 | 61 | 28.17 | 446 | 3348605 | 34 | 3 | 2999 | 3037 | 97.26 | positive |
| 43 | PH091 | Individual 1 | 2013 | London | FP | 2 | MiSeq | 101 | 881394 | 53 | 28.17 | 491 | 3337668 | 35 | 4 | 2968 | 3008 | 97.25 | positive |
| 44 | PH024 | Individual 3 | 2014 | North east | FP | 3 | HiSeq | 151 | 3024896 | 274 | 28.03 | 28 | 3331444 | 90 | 12 | 3016 | 3119 | 97.23 | positive |
| 45 | PH025 | Individual 4 | 2014 | North east | FP | 3 | HiSeq | 151 | 2413851 | 219 | 28 | 34 | 3320821 | 72 | 7 | 3002 | 3082 | 97.23 | positive |
| 46 | PH023 | Individual 2 | 2014 | North east | FP | 3 | HiSeq | 151 | 2909655 | 264 | 28.01 | 35 | 3325484 | 88 | 7 | 3002 | 3098 | 97.23 | positive |
| 47 | PH022 | Individual 1 | 2014 | North east | FP | 3 | HiSeq | 151 | 2319050 | 205 | 28.01 | 36 | 3401212 | 74 | 12 | 3077 | 3164 | 97.24 | positive |
| 48 | PH026 | Cooked sliced beef | 2014 | North east | FP | 3 | HiSeq | 151 | 2769966 | 250 | 28.07 | 37 | 3335204 | 91 | 12 | 3009 | 3113 | 97.24 | positive |
| 49 | PH028 | Individual 4 | 2014 | London | FP | 4 | HiSeq | 151 | 3031314 | 305 | 27.96 | 82 | 2994736 | 75 | 10 | 2731 | 2817 | 99.64 | positive |
| 50 | PH029 | Individual 5 | 2014 | London | FP | 4 | HiSeq | 151 | 3106630 | 317 | 27.92 | 97 | 2954967 | 87 | 12 | 2694 | 2794 | 99.65 | positive |
| 51 | PH104 | Individual 1 | 2014 | London | FP | 4 | MiSeq | 101 | 1072977 | 73 | 27.9 | 214 | 2929035 | 43 | 4 | 2668 | 2716 | 99.64 | positive |
| 52 | PH107 | Individual 6 | 2014 | London | FP | 4 | MiSeq | 101 | 877265 | 60 | 27.91 | 246 | 2914363 | 38 | 5 | 2650 | 2694 | 99.66 | positive |
| 53 | PH105 | Individual 3 | 2014 | North East | FP | 4 | MiSeq | 101 | 1022558 | 70 | 27.93 | 379 | 2944409 | 57 | 7 | 2672 | 2737 | 99.65 | positive |
| 54 | PH115 | Individual 2 | 2014 | North east | FP | 5 | MiSeq | 101 | 1537158 | 104 | 27.87 | 317 | 2977232 | 68 | 6 | 2694 | 2769 | 99.51 | positive |
| 55 | PH118 | Individual 2 | 2015 | North East | FP | 6 | MiSeq | 101 | 1632880 | 100 | 28.01 | 214 | 3282569 | 62 | 7 | 2917 | 2987 | 97.21 | positive |
| 56 | PH119 | Individual 3 | 2015 | North East | FP | 6 | MiSeq | 101 | 1287449 | 79 | 28 | 215 | 3289115 | 66 | 6 | 2913 | 2986 | 97.20 | positive |
| 57 | PH117 | Individual 1 | 2015 | North East | FP | 6 | MiSeq | 101 | 1569491 | 96 | 28.02 | 246 | 3285816 | 74 | 10 | 2917 | 3002 | 97.22 | positive |
| 58 | PH039 | Individual 4 | 2015 | North East | FP | 7 | HiSeq | 151 | 2709205 | 238 | 28.13 | 23 | 3432249 | 86 | 11 | 3094 | 3192 | 97.22 | positive |
| 59 | PH040 | Individual 5 | 2015 | North East | FP | 7 | HiSeq | 151 | 2211747 | 194 | 28.12 | 25 | 3426487 | 87 | 11 | 3089 | 3188 | 97.21 | positive |
| 60 | PH037 | Individual 3 | 2015 | North East | FP | 7 | HiSeq | 151 | 2113659 | 186 | 28.12 | 26 | 3426315 | 71 | 10 | 3088 | 3170 | 97.21 | positive |
| 61 | PH044 | Individual 8 | 2015 | North East | FP | 7 | HiSeq | 151 | 1993382 | 175 | 28.12 | 29 | 3426828 | 83 | 9 | 3088 | 3181 | 97.21 | positive |
| 62 | PH036 | Individual 2 | 2015 | North East | FP | 7 | HiSeq | 151 | 2238843 | 197 | 28.14 | 30 | 3430269 | 88 | 13 | 3090 | 3192 | 97.21 | positive |
| 63 | PH041 | Individual 6 | 2015 | North East | FP | 7 | HiSeq | 151 | 1997998 | 175 | 28.13 | 31 | 3433511 | 88 | 12 | 3099 | 3200 | 97.21 | positive |
| 64 | PH035 | Individual 1 | 2015 | North East | FP | 7 | HiSeq | 151 | 1947365 | 171 | 28.11 | 32 | 3422738 | 86 | 10 | 3081 | 3178 | 97.21 | positive |
| 65 | PH045 | Individual 8 | 2015 | North East | FP | 7 | HiSeq | 151 | 2313617 | 218 | 28.07 | 32 | 3191860 | 90 | 9 | 2864 | 2964 | 97.23 | negative |
| 66 | PH042 | Individual 11 | 2015 | North East | FP | 7 | HiSeq | 151 | 1766545 | 155 | 28.12 | 34 | 3427997 | 87 | 13 | 3087 | 3188 | 97.21 | positive |
| 67 | PH048 | Individual 12 | 2015 | North East | FP | 7 | HiSeq | 151 | 2875352 | 253 | 28.13 | 34 | 3428504 | 92 | 8 | 3087 | 3188 | 97.21 | positive |
| 68 | PH046 | Individual 9 | 2015 | North East | FP | 7 | HiSeq | 151 | 2091933 | 184 | 28.11 | 35 | 3425020 | 93 | 9 | 3080 | 3183 | 97.20 | positive |
| 69 | PH043 | Individual 7 | 2015 | North East | FP | 7 | HiSeq | 151 | 2248776 | 198 | 28.1 | 43 | 3421670 | 69 | 8 | 3081 | 3159 | 97.20 | positive |
| 70 | PH047 | Individual 10 | 2015 | North East | FP | 7 | HiSeq | 151 | 2214491 | 195 | 28.11 | 43 | 3422937 | 71 | 9 | 3084 | 3165 | 97.21 | positive |
| 71 | PH154 | Individual | 2015 | North East | FP | 7 | MiSeq | 101 | 1800754 | 106 | 28.11 | 177 | 3413080 | 73 | 8 | 3065 | 3147 | 97.22 | positive |
| 72 | PH155 | Individual 8 | 2015 | North East | FP | 7 | MiSeq | 101 | 1230813 | 72 | 28.11 | 209 | 3411087 | 58 | 7 | 3069 | 3135 | 97.23 | positive |
| 73 | PH156 | Individual 8 | 2015 | North East | FP | 7 | MiSeq | 101 | 1206348 | 76 | 28.07 | 239 | 3172681 | 51 | 5 | 2846 | 2903 | 97.25 | negative |
| 74 | PH050 | Individual 1 | 2015 | South West | FP | 8 | HiSeq | 151 | 2468557 | 224 | 28.13 | 59 | 3315627 | 93 | 12 | 2944 | 3050 | 97.21 | positive |
| 75 | PH120 | Individual 2 | 2015 | South West | FP | 8 | MiSeq | 101 | 1102939 | 68 | 28.05 | 185 | 3274008 | 40 | 4 | 2894 | 2939 | 97.22 | positive |
| 76 | PH121 | Lamb | 2015 | South West | FP | 8 | MiSeq | 101 | 1382732 | 86 | 28.08 | 289 | 3233465 | 38 | 8 | 2854 | 2901 | 97.22 | positive |
| 77 | PH122 | Lamb | 2015 | South West | FP | 8 | MiSeq | 101 | 1008241 | 62 | 28.11 | 371 | 3282469 | 70 | 7 | 2901 | 2979 | 97.24 | positive |
| 78 | PH055 | Individual 3 | 2015 | North east | FP | 9 | HiSeq | 151 | 3085261 | 270 | 28 | 27 | 3441127 | 76 | 9 | 3130 | 3216 | 97.28 | positive |
| 79 | PH054 | Chicken curry | 2015 | North east | FP | 9 | HiSeq | 151 | 2270852 | 197 | 28.03 | 57 | 3476853 | 87 | 12 | 3153 | 3253 | 97.27 | positive |
| 80 | PH124 | Individual 2 | 2015 | North east | FP | 9 | MiSeq | 101 | 1394099 | 82 | 27.99 | 132 | 3431905 | 46 | 5 | 3120 | 3172 | 97.29 | positive |
| 81 | PH123 | Individual 1 | 2015 | North east | FP | 9 | MiSeq | 101 | 1445464 | 85 | 28 | 156 | 3429751 | 67 | 5 | 3113 | 3186 | 97.29 | positive |
| 82 | PH057 | Individual 2 | 2015 | North east | FP | 10 | HiSeq | 151 | 2899190 | 257 | 28.12 | 18 | 3398027 | 92 | 10 | 3037 | 3140 | 97.20 | positive |
| 83 | PH066 | 3 birds roast | 2015 | North east | FP | 10 | HiSeq | 151 | 2252903 | 200 | 28.13 | 18 | 3398355 | 90 | 11 | 3037 | 3139 | 97.20 | positive |
| 84 | PH063 | Individual 8 | 2015 | North east | FP | 10 | HiSeq | 151 | 2652135 | 235 | 28.11 | 19 | 3395429 | 75 | 10 | 3037 | 3123 | 97.19 | positive |
| 85 | PH060 | Individual 5 | 2015 | North east | FP | 10 | HiSeq | 151 | 2421609 | 215 | 28.13 | 20 | 3399028 | 89 | 12 | 3038 | 3140 | 97.19 | positive |
| 86 | PH061 | Individual 6 | 2015 | North east | FP | 10 | HiSeq | 151 | 2774093 | 246 | 28.12 | 20 | 3396723 | 89 | 10 | 3037 | 3137 | 97.19 | positive |
| 87 | PH065 | Individual 10 | 2015 | North east | FP | 10 | HiSeq | 151 | 2302879 | 204 | 28.1 | 20 | 3392964 | 93 | 8 | 3037 | 3139 | 97.19 | positive |
| 88 | PH062 | Individual 7 | 2015 | North east | FP | 10 | HiSeq | 151 | 2623765 | 233 | 28.12 | 21 | 3397544 | 87 | 9 | 3037 | 3134 | 97.20 | positive |
| 89 | PH067 | Individual 11 | 2015 | North east | FP | 10 | HiSeq | 151 | 2428595 | 216 | 28.1 | 21 | 3392663 | 75 | 8 | 3036 | 3120 | 97.18 | positive |
| 90 | PH058 | Individual 3 | 2015 | North east | FP | 10 | HiSeq | 151 | 2571555 | 228 | 28.12 | 22 | 3396864 | 83 | 12 | 3036 | 3132 | 97.19 | positive |
| 91 | PH056 | Individual 1 | 2015 | North east | FP | 10 | HiSeq | 151 | 2422080 | 215 | 28.11 | 23 | 3395441 | 80 | 10 | 3037 | 3128 | 97.19 | positive |
| 92 | PH068 | Individual 12 | 2015 | North east | FP | 10 | HiSeq | 151 | 2060812 | 183 | 28.13 | 23 | 3399957 | 87 | 10 | 3037 | 3135 | 97.20 | positive |
| 93 | PH059 | Individual 4 | 2015 | North east | FP | 10 | HiSeq | 151 | 2864609 | 254 | 28.11 | 24 | 3395953 | 89 | 9 | 3037 | 3136 | 97.19 | positive |
| 94 | PH127 | Individual 3 | 2016 | Newcastle | FP | 11 | MiSeq | 101 | 1014192 | 70 | 27.93 | 262 | 2924057 | 50 | 7 | 2649 | 2707 | 99.47 | positive |
| 95 | PH125 | Individual 1 | 2015 | Newcastle | FP | 11 | MiSeq | 101 | 1011178 | 59 | 28.09 | 363 | 3406444 | 27 | 3 | 3089 | 3120 | 97.31 | positive |
| 96 | PH126 | Individual 2 | 2016 | Newcastle | FP | 11 | MiSeq | 101 | 1153871 | 68 | 28.09 | 418 | 3408431 | 32 | 4 | 3077 | 3114 | 97.31 | positive |
| 97 | PH134 | Cooked turkey | 2016 | West Midlands | FP | 12 | MiSeq | 101 | 1617315 | 94 | 28.08 | 192 | 3446865 | 76 | 8 | 3077 | 3162 | 97.17 | positive |
| 98 | PH130 | Individual 3 | 2016 | West Midlands | FP | 12 | MiSeq | 101 | 1374433 | 80 | 28.09 | 250 | 3446961 | 68 | 7 | 3078 | 3154 | 97.16 | positive |
| 99 | PH129 | Individual 2 | 2016 | West Midlands | FP | 12 | MiSeq | 101 | 1599870 | 93 | 28.09 | 266 | 3441457 | 57 | 6 | 3070 | 3134 | 97.17 | positive |
| 100 | PH133 | Cooked meat | 2016 | West Midlands | FP | 12 | MiSeq | 101 | 1833668 | 107 | 28.11 | 347 | 3432746 | 54 | 6 | 3055 | 3116 | 97.18 | positive |
| 101 | PH128 | Individual 1 | 2016 | West Midlands | FP | 12 | MiSeq | 101 | 1316831 | 77 | 28.09 | 352 | 3436609 | 45 | 8 | 3064 | 3118 | 97.20 | positive |
| 102 | PH132 | Individual 6 | 2016 | West Midlands | FP | 12 | MiSeq | 101 | 1569588 | 92 | 28.14 | 466 | 3418685 | 57 | 8 | 3029 | 3095 | 97.21 | positive |
| 103 | PH136 | Individual 2 | 2016 | London | FP | 13 | MiSeq | 101 | 1317728 | 91 | 27.91 | 312 | 2905546 | 45 | 5 | 2638 | 2689 | 99.40 | positive |
| 104 | PH135 | Individual 1 | 2016 | London | FP | 13 | MiSeq | 101 | 1152631 | 79 | 27.9 | 356 | 2912738 | 40 | 6 | 2644 | 2691 | 99.40 | positive |
| 105 | PH081 | Individual 3 | 2017 | North West | FP | 14 | HiSeq | 151 | 2085111 | 211 | 27.89 | 67 | 2984289 | 91 | 11 | 2729 | 2832 | 99.47 | positive |
| 106 | PH078 | Individual 1 | 2017 | North West | FP | 14 | HiSeq | 151 | 2152453 | 217 | 27.92 | 70 | 2987740 | 89 | 13 | 2727 | 2830 | 99.47 | positive |
| 107 | PH080 | Individual 2 | 2017 | North West | FP | 14 | HiSeq | 151 | 2501153 | 253 | 27.9 | 73 | 2983246 | 89 | 11 | 2718 | 2819 | 99.47 | positive |
| 108 | PH143 | Individual 1 | 2016 | South West | FP | sporadic | MiSeq | 101 | 1333457 | 76 | 28.05 | 256 | 3528864 | 56 | 6 | 3176 | 3239 | 97.24 | positive |
| 109 | PH147 | Individual 1 | 2013 | North East | FP | sporadic | MiSeq | 101 | 1204345 | 74 | 28.03 | 351 | 3267085 | 53 | 6 | 2918 | 2978 | 97.28 | positive |
