## Supplementary Table 9 for "Phylogenomic analysis of *Clostridium perfringens* identifies isogenic strains in gastroenteritis outbreaks, and novel virulence-related features"

**Table S9. Similarity comparisons of predicted plasmids with reference plasmids**

Predicted plasmids were compared with reference plasmids pCPF5603 and pCPF4969.K-mer identity (based on FASTQ) determined by PlasmidSeeker; BLASTN sequence similarity and coverage determined by ABRicate v0.5. Genome features of plasmids were extracted from Prokka annotations.

| Reference plasmid | Predicted Plasmid | Genome features | | | K-mer identity (%)  (Read-based) | Similarity by BLASTN  (Assembly-based) | |
| --- | --- | --- | --- | --- | --- | --- | --- |
|  |  | Size (bp) | GC (%) | CDS |  | Identity (%) | Coverage (%) |
| **pCPF5603** | pPH077 | 75,398 | 25.54 | 87 | 99.64% | 99.98 | 100.00 |
|  | pPH032 | 75,402 | 25.53 | 87 | 99.58% | 99.98 | 100.00 |
|  | pPH031 | 75,401 | 25.54 | 87 | 99.63% | 99.97 | 100.00 |
| **pCPF4969** | pPH062 | 70,638 | 26.59 | 74 | 99.66% | 99.98 | 100.00 |
|  | pPH041 | 70,811 | 26.57 | 75 | 99.53% | 99.97 | 100.00 |
|  | pPH050 | 70,624 | 26.57 | 75 | 99.40% | 99.98 | 100.00 |
