## Supplementary Table 10 for "Phylogenomic analysis of *Clostridium perfringens* identifies isogenic strains in gastroenteritis outbreaks, and novel virulence-related features"

**Table S10. Genome comparison of isolates within FP lineage I**

Genome sizes were based on genome assemblies of isolates. ANI (%) of each genome was compared with reference genome NCTC8239.

| Isolate | FP Lineage | Genome Size (bp) | GC (%) | ANI (%) |
| --- | --- | --- | --- | --- |
| NCTC8239 | I | 3,008,497 | 28.19 | 100.00 |
| PH004 | I | 2,923,443 | 27.87 | 99.42 |
| PH005 | I | 2,934,006 | 27.93 | 99.43 |
| PH006 | I | 2,928,275 | 27.89 | 99.42 |
| PH007 | I | 2,946,980 | 27.89 | 99.42 |
| PH135 | I | 2,912,738 | 27.90 | 99.40 |
| PH136 | I | 2,905,546 | 27.91 | 99.40 |
| PH115 | I | 2,977,232 | 27.87 | 99.51 |
| PH127 | I | 2,924,057 | 27.93 | 99.47 |
| PH080 | I | 2,983,246 | 27.90 | 99.47 |
| PH078 | I | 2,987,740 | 27.92 | 99.47 |
| PH081 | I | 2,984,289 | 27.89 | 99.47 |
| PH028 | I | 2,994,736 | 27.96 | 99.64 |
| PH029 | I | 2,954,967 | 27.92 | 99.65 |
| PH104 | I | 2,929,035 | 27.90 | 99.64 |
| PH107 | I | 2,914,363 | 27.91 | 99.66 |
| PH105 | I | 2,944,409 | 27.93 | 99.65 |
| **Stats (mean ± SD)** | | **2.95 ± 0.03 Mb** | **27.92 ± 0.07** | **99.51 ± 0.10** |
