## Supplementary Table 3 for "Phylogenomic analysis of *Clostridium perfringens* identifies isogenic strains in gastroenteritis outbreaks, and novel virulence-related features"

**Table S3.** RefSeq genomes used to generate *Clostridium* genus-specific database for genome annotation (Prokka). *Recently suggested to be moved to genus *Clostridioides*. ^#^Recently suggested to be classified as genus *Paeniclostridium*.

|  | **RefSeq Genome Species** | **Strain** | **NCBI Accession** |
| --- | --- | --- | --- |
| 1 | *C. perfringens* | 1207_CPER | SAMN03197169 |
| 2 | *C. perfringens* | ATCC13124 | SAMN02604008 |
| 3 | *C. perfringens* | B_ATCC3626 | SAMN02436295 |
| 4 | *C. perfringens* | CBA7123 | SAMD00057672 |
| 5 | *C. perfringens* | C_JGS1495 | SAMN02436294 |
| 6 | *C. perfringens* | CP15 | SAMN06252080 |
| 7 | *C. perfringens* | CPE_F4969 | SAMN02436168 |
| 8 | *C. perfringens* | D_JGS1721 | SAMN02436277 |
| 9 | *C. perfringens* | E_JGS1987 | SAMN02436167 |
| 10 | *C. perfringens* | FORC003 | SAMN03140316 |
| 11 | *C. perfringens* | FORC025 | SAMN04209542 |
| 12 | *C. perfringens* | JFP718 | SAMN05323879 |
| 13 | *C. perfringens* | JFP836 | SAMN05323894 |
| 14 | *C. perfringens* | JJC | SAMN02317206 |
| 15 | *C. perfringens* | JP55 | SAMN03372134 |
| 16 | *C. perfringens* | JP838 | SAMN03377063 |
| 17 | *C. perfringens* | LLY_N11 | SAMN07624709 |
| 18 | *C. perfringens* | NCTC8239 | SAMN02436239 |
| 19 | *C. perfringens* | 13 | SAMD00061119 |
| 20 | *C. perfringens* | 2789STDY5608889 | SAMEA3545297 |
| 21 | *C. acetobutylicum* | ATCC824 | SAMN02603243 |
| 22 | *C. baratti* | Sullivan | SAMN03222823 |
| 23 | *C. botulinum* | ATCC3502 | SAMEA1705919 |
| 24 | *C. butyricum* | KNU-L09 | SAMN04293668 |
| 25 | *C. chauvoei* | JF4335 | SAMEA102059668 |
| 26 | *C. difficile** | 630 | SAMEA1705932 |
| 27 | *C. difficile ** | BR81 | SAMN06286529 |
| 28 | *C. difficile** | E15 | SAMEA3138880 |
| 29 | *C. difficile** | M120 | SAMEA3138368 |
| 30 | *C. novyi* | NT | SAMN02603464 |
| 31 | *C. paraputrificum* | LH025 | SAMEA104287982 |
| 32 | *C. pasteurianum* | DSM525 | SAMN02990135 |
| 33 | *C. sordellii^#^* | AM370 | SAMN04263499 |
| 34 | *C. tertium* | LH009 | SAMEA104287969 |
| 35 | *C. tetani* | E88 | SAMN02603289 |
