## Supplementary Table 4 for "Phylogenomic analysis of *Clostridium perfringens* identifies isogenic strains in gastroenteritis outbreaks, and novel virulence-related features"

**Table S4.** *C. perfringens* plasmid sequences from NCBI RefSeq database included in PlasmidSeeker database.

|  | Plasmid sequence names |
| --- | --- |
| 1 | *Clostridium perfringens* plasmid pCP-TS1 DNA, complete sequence, strain: TS1 |
| 2 | *Clostridium perfringens* plasmid pCP8533etx, complete sequence |
| 3 | *Clostridium perfringens* plasmid pCP-OS1 DNA, complete sequence, strain: OS1 |
| 4 | *Clostridium perfringens* plasmid pCPPB-1, complete sequence |
| 5 | *Clostridium perfringens* plasmid pCW3, complete sequence |
| 6 | *Clostridium perfringens* plasmid pIP404, complete sequence |
| 7 | *Clostridium perfringens* plasmid pJIR3536, complete sequence |
| 8 | *Clostridium perfringens* plasmid pCPF5603, complete sequence |
| 9 | *Clostridium perfringens* plasmid pBCNF5603 DNA, complete sequence |
| 10 | *Clostridium perfringens* plasmid pCP8533S12 DNA, complete sequence |
| 11 | *Clostridium perfringens* strain FORC_003 plasmid pFORC3, complete sequence |
| 12 | *Clostridium perfringens* strain JP55 plasmid pJFP55F, complete sequence |
| 13 | *Clostridium perfringens* strain JP55 plasmid pJFP55G |
| 14 | *Clostridium perfringens* strain JP55 plasmid pJFP55H, complete sequence |
| 15 | *Clostridium perfringens* strain JP55 plasmid pJFP55J, complete sequence |
| 16 | *Clostridium perfringens* strain JP55 plasmid pJFP55K, complete sequence |
| 17 | *Clostridium perfringens* strain JP838 plasmid pJFP838E, complete sequence |
| 18 | *Clostridium perfringens* strain JP838 plasmid pJFP838D, complete sequence |
| 19 | *Clostridium perfringens* strain JP838 plasmid pJFP838C, complete sequence |
| 20 | *Clostridium perfringens* strain JP838 plasmid pJFP838A, complete sequence |
| 21 | *Clostridium perfringens* str. 13 plasmid pCP13 DNA, complete sequence |
| 22 | *Clostridium perfringens* CPE str. F4969 plasmid pCPF4969, complete sequence |
| 23 | *Clostridium perfringens* plasmid pJIR3844, complete sequence |
| 24 | *Clostridium perfringens* plasmid pJIR3843, complete sequence |
| 25 | *Clostridium perfringens* plasmid pJIR3537, complete sequence |
| 26 | *Clostridium perfringens* plasmid pCpb2-CP1, complete sequence |
| 27 | *Clostridium perfringens* plasmid pNetB-NE10, complete sequence |
| 28 | *Clostridium perfringens* strain Del1 plasmid pDel1_1, complete sequence |
| 29 | *Clostridium perfringens* strain Del1 plasmid pDel1_2, complete sequence |
| 30 | *Clostridium perfringens* strain Del1 plasmid pDle1_3, complete sequence |
| 31 | *Clostridium perfringens* strain Del1 plasmid pDel1_4, complete sequence |
| 32 | *Clostridium perfringens* strain CP15 plasmid pCP15_1, complete sequence |
| 33 | *Clostridium perfringens* strain CP15 plasmid pCP15_2, complete sequence |
| 34 | *Clostridium perfringens* strain CP15 plasmid pCP15_3, complete sequence |
| 35 | *Clostridium perfringens* strain CP15 plasmid pCP15_4, complete sequence |
