## Supplementary Table 5 for "Phylogenomic analysis of *Clostridium perfringens* identifies isogenic strains in gastroenteritis outbreaks, and novel virulence-related features"

**Table S5.** Plasmids and related sequences predicted in all 109 *C. perfringens* isolates in this study.

|  | **Isolate information** | | **Plasmid-related sequences** | | | **Predicted plasmid 1** | | | **Predicted plasmid 2** | | | **Predicted plasmid 3** | | |
| --- | --- | --- | --- | --- | --- | --- | --- | --- | --- | --- | --- | --- | --- | --- |
|  | Isolate | Cohort | IS1151 | IS1470-like | tcp locus gene | Plasmid | K-mer % | Copy number | Plasmid | K-mer % | Copy number | Plasmid | K-mer % | Copy number |
| 1 | PH015 | CH | 1 | 0 | 8 | pCPF5603 | 99.55% | 1.44 |  |  |  |  |  |  |
| 2 | PH016 | CH | 1 | 0 | 8 | pCPF5603 | 99.55% | 2.06 |  |  |  |  |  |  |
| 3 | PH017 | CH | 0 | 0 | 8 | pFORC3 | 81.80% | 2.64 |  |  |  |  |  |  |
| 4 | PH018 | CH | 1 | 0 | 8 | pCPF5603 | 99.64% | 1.26 |  |  |  |  |  |  |
| 5 | PH019 | CH | 1 | 0 | 8 | pCPF5603 | 99.65% | 1.66 |  |  |  |  |  |  |
| 6 | PH020 | CH | 1 | 0 | 8 | pCPF5603 | 99.58% | 2.56 | pBCNF5603 | 95.71% | 1.98 |  |  |  |
| 7 | PH021 | CH | 1 | 0 | 8 | pCPF5603 | 99.59% | 1.79 | pBCNF5603 | 96.04% | 1.88 |  |  |  |
| 8 | PH030 | CH | 1 | 0 | 8 | pCPF5603 | 99.63% | 1.52 |  |  |  |  |  |  |
| 9 | PH031 | CH | 1 | 0 | 8 | pCPF5603 | 99.63% | 1.39 |  |  |  |  |  |  |
| 10 | PH032 | CH | 1 | 0 | 8 | pCPF5603 | 99.58% | 1.89 |  |  |  |  |  |  |
| 11 | PH033 | CH | 1 | 0 | 8 | pCPF5603 | 99.55% | 1.49 |  |  |  |  |  |  |
| 12 | PH051 | CH | 1 | 0 | 8 | pCPF5603 | 99.60% | 1.96 |  |  |  |  |  |  |
| 13 | PH052 | CH | 1 | 0 | 8 | pCPF5603 | 99.66% | 1.77 |  |  |  |  |  |  |
| 14 | PH053 | CH | 1 | 0 | 8 | pCPF5603 | 99.63% | 1.76 |  |  |  |  |  |  |
| 15 | PH072 | CH | 1 | 0 | 8 | pCPF5603 | 89.17% | 1.07 | pFORC3 | 83.40% | 1.09 |  |  |  |
| 16 | PH073 | CH | 1 | 0 | 8 | pCPF5603 | 99.62% | 1.84 |  |  |  |  |  |  |
| 17 | PH074 | CH | 1 | 0 | 8 | pCPF5603 | 99.58% | 1.43 |  |  |  |  |  |  |
| 18 | PH075 | CH | 1 | 0 | 8 | pCPF5603 | 99.58% | 1.57 |  |  |  |  |  |  |
| 19 | PH076 | CH | 1 | 0 | 8 | pCPF5603 | 99.66% | 1.5 |  |  |  |  |  |  |
| 20 | PH077 | CH | 1 | 0 | 8 | pCPF5603 | 99.64% | 1.7 |  |  |  |  |  |  |
| 21 | PH079 | CH | 1 | 0 | 8 | pCPF5603 | 99.62% | 1.59 |  |  |  |  |  |  |
| 22 | PH082 | CH | 1 | 0 | 8 | pCPF5603 | 99.66% | 1.71 |  |  |  |  |  |  |
| 23 | PH083 | CH | 1 | 0 | 8 | pCPF5603 | 99.66% | 2.76 |  |  |  |  |  |  |
| 24 | PH084 | CH | 1 | 0 | 8 | pCPF5603 | 99.66% | 1.7 |  |  |  |  |  |  |
| 25 | PH085 | CH | 1 | 0 | 8 | pCPF5603 | 99.66% | 1.61 |  |  |  |  |  |  |
| 26 | PH096 | CH | 1 | 0 | 8 | pCPF5603 | 99.66% | 0.88 |  |  |  |  |  |  |
| 27 | PH097 | CH | 1 | 0 | 8 | pCPF5603 | 88.39% | 0.79 |  |  |  |  |  |  |
| 28 | PH098 | CH | 1 | 0 | 8 | pCPF5603 | 93.41% | 0.8 |  |  |  |  |  |  |
| 29 | PH100 | CH | 1 | 0 | 8 | pCPF5603 | 80.18% | 0.83 |  |  |  |  |  |  |
| 30 | PH101 | CH | 1 | 0 | 8 | pCPF5603 | 96.18% | 0.78 |  |  |  |  |  |  |
| 31 | PH102 | CH | 1 | 0 | 8 | pCPF5603 | 91.15% | 0.79 |  |  |  |  |  |  |
| 32 | PH103 | CH | 1 | 0 | 8 | pCPF5603 | 93.91% | 0.86 |  |  |  |  |  |  |
| 33 | PH138 | CH | 1 | 0 | 8 | pCPF5603 | 84.57% | 0.79 | pBCNF5603 | 82.38% | 0.75 |  |  |  |
| 34 | PH139 | CH | 1 | 0 | 8 | pCPF5603 | 91.75% | 0.96 | pBCNF5603 | 91.79% | 1.26 |  |  |  |
| 35 | PH140 | CH | 1 | 0 | 8 | pCPF5603 | 86.62% | 0.88 | pBCNF5603 | 84.18% | 0.8 |  |  |  |
| 36 | PH004 | FP | 0 | 0 | 0 | 0 |  |  |  |  |  |  |  |  |
| 37 | PH005 | FP | 0 | 0 | 0 | 0 |  |  |  |  |  |  |  |  |
| 38 | PH006 | FP | 0 | 0 | 0 | 0 |  |  |  |  |  |  |  |  |
| 39 | PH007 | FP | 0 | 0 | 0 | 0 |  |  |  |  |  |  |  |  |
| 40 | PH014 | FP | 0 | 1 | 10 | pCPF4969 | 98.99% | 2.84 | pCW3 | 98.46% | 3.17 | pDel1_4 | 93.68% | 3.17 |
| 41 | PH022 | FP | 0 | 1 | 10 | pCPF4969 | 99.20% | 2.53 |  |  |  |  |  |  |
| 42 | PH023 | FP | 0 | 1 | 10 | pCPF4969 | 99.25% | 2.17 |  |  |  |  |  |  |
| 43 | PH024 | FP | 0 | 1 | 10 | pCPF4969 | 99.38% | 3.11 |  |  |  |  |  |  |
| 44 | PH025 | FP | 0 | 1 | 10 | pCPF4969 | 99.20% | 2.45 |  |  |  |  |  |  |
| 45 | PH026 | FP | 0 | 1 | 10 | pCPF4969 | 99.45% | 1.84 |  |  |  |  |  |  |
| 46 | PH028 | FP | 0 | 0 | 0 | 0 |  |  |  |  |  |  |  |  |
| 47 | PH029 | FP | 0 | 0 | 0 | 0 |  |  |  |  |  |  |  |  |
| 48 | PH035 | FP | 0 | 1 | 10 | pCPF4969 | 99.55% | 2.59 |  |  |  |  |  |  |
| 49 | PH036 | FP | 0 | 1 | 10 | pCPF4969 | 99.56% | 2.45 |  |  |  |  |  |  |
| 50 | PH037 | FP | 0 | 1 | 10 | pCPF4969 | 99.59% | 2.21 |  |  |  |  |  |  |
| 51 | PH039 | FP | 0 | 1 | 10 | pCPF4969 | 99.55% | 2.31 |  |  |  |  |  |  |
| 52 | PH040 | FP | 0 | 1 | 10 | pCPF4969 | 99.55% | 4.76 |  |  |  |  |  |  |
| 53 | PH041 | FP | 0 | 1 | 10 | pCPF4969 | 99.53% | 1.97 |  |  |  |  |  |  |
| 54 | PH042 | FP | 0 | 1 | 10 | pCPF4969 | 99.52% | 3.37 |  |  |  |  |  |  |
| 55 | PH043 | FP | 0 | 1 | 10 | pCPF4969 | 99.53% | 2.69 |  |  |  |  |  |  |
| 56 | PH044 | FP | 0 | 1 | 10 | pCPF4969 | 99.55% | 3.73 |  |  |  |  |  |  |
| 57 | PH045 | FP | 0 | 0 | 0 |  |  |  |  |  |  |  |  |  |
| 58 | PH046 | FP | 0 | 1 | 10 | pCPF4969 | 99.52% | 2.76 |  |  |  |  |  |  |
| 59 | PH047 | FP | 0 | 1 | 10 | pCPF4969 | 99.60% | 2.61 |  |  |  |  |  |  |
| 60 | PH048 | FP | 0 | 1 | 10 | pCPF4969 | 99.55% | 2.77 |  |  |  |  |  |  |
| 61 | PH050 | FP | 0 | 1 | 10 | pCPF4969 | 99.40% | 1.49 |  |  |  |  |  |  |
| 62 | PH054 | FP | 1 | 0 | 8 | pCPF5603 | 99.61% | 2.24 |  |  |  |  |  |  |
| 63 | PH055 | FP | 1 | 0 | 8 | pCPF5603 | 99.64% | 1.99 |  |  |  |  |  |  |
| 64 | PH056 | FP | 0 | 1 | 10 | pCPF4969 | 99.68% | 4.99 |  |  |  |  |  |  |
| 65 | PH057 | FP | 0 | 1 | 10 | pCPF4969 | 99.65% | 2.42 |  |  |  |  |  |  |
| 66 | PH058 | FP | 0 | 1 | 10 | pCPF4969 | 99.69% | 2.25 |  |  |  |  |  |  |
| 67 | PH059 | FP | 0 | 1 | 10 | pCPF4969 | 99.65% | 3.21 |  |  |  |  |  |  |
| 68 | PH060 | FP | 0 | 1 | 10 | pCPF4969 | 99.63% | 2.28 |  |  |  |  |  |  |
| 69 | PH061 | FP | 0 | 1 | 10 | pCPF4969 | 99.63% | 2.28 |  |  |  |  |  |  |
| 70 | PH062 | FP | 0 | 1 | 10 | pCPF4969 | 99.66% | 2 |  |  |  |  |  |  |
| 71 | PH063 | FP | 0 | 1 | 10 | pCPF4969 | 99.67% | 2.09 |  |  |  |  |  |  |
| 72 | PH065 | FP | 0 | 1 | 10 | pCPF4969 | 99.63% | 1.57 |  |  |  |  |  |  |
| 73 | PH066 | FP | 0 | 1 | 10 | pCPF4969 | 99.63% | 2.16 |  |  |  |  |  |  |
| 74 | PH067 | FP | 0 | 1 | 10 | pCPF4969 | 99.64% | 1.88 |  |  |  |  |  |  |
| 75 | PH068 | FP | 0 | 1 | 10 | pCPF4969 | 99.63% | 2.2 |  |  |  |  |  |  |
| 76 | PH078 | FP | 0 | 0 | 0 |  |  |  |  |  |  |  |  |  |
| 77 | PH080 | FP | 0 | 0 | 0 |  |  |  |  |  |  |  |  |  |
| 78 | PH081 | FP | 0 | 0 | 0 |  |  |  |  |  |  |  |  |  |
| 79 | PH091 | FP | 0 | 1 | 10 | pCPF4969 | 94.39% | 1.69 |  |  |  |  |  |  |
| 80 | PH092 | FP | 0 | 1 | 10 | pCPF4969 | 88.48% | 2.46 | pCW3 | 89.49% | 2.92 |  |  |  |
| 81 | PH095 | FP | 0 | 1 | 10 | pCPF4969 | 85.19% | 1.89 | pCW3 | 87.20% | 2.53 |  |  |  |
| 82 | PH104 | FP | 0 | 0 | 0 |  |  |  |  |  |  |  |  |  |
| 83 | PH105 | FP | 0 | 0 | 0 |  |  |  |  |  |  |  |  |  |
| 84 | PH107 | FP | 0 | 0 | 0 |  |  |  |  |  |  |  |  |  |
| 85 | PH115 | FP | 0 | 0 | 0 |  |  |  |  |  |  |  |  |  |
| 86 | PH117 | FP | 0 | 1 | 10 | pCPF4969 | 98.34% | 1.38 |  |  |  |  |  |  |
| 87 | PH118 | FP | 0 | 1 | 10 | pCPF4969 | 94.66% | 1.31 |  |  |  |  |  |  |
| 88 | PH119 | FP | 0 | 1 | 10 | pCPF4969 | 95.07% | 1.44 |  |  |  |  |  |  |
| 89 | PH120 | FP | 0 | 1 | 10 | pCPF4969 | 98.09% | 1.26 |  |  |  |  |  |  |
| 90 | PH121 | FP | 0 | 1 | 10 | pCPF4969 | 95.03% | 1.15 |  |  |  |  |  |  |
| 91 | PH122 | FP | 0 | 1 | 10 | pCPF4969 | 96.19% | 1.19 |  |  |  |  |  |  |
| 92 | PH123 | FP | 1 | 0 | 8 | pCPF5603 | 93.98% | 0.89 |  |  |  |  |  |  |
| 93 | PH124 | FP | 1 | 0 | 8 | pCPF5603 | 95.64% | 1 |  |  |  |  |  |  |
| 94 | PH125 | FP | 1 | 0 | 8 | pBCNF5603 | 82.68% | 0.75 |  |  |  |  |  |  |
| 95 | PH126 | FP | 1 | 0 | 8 | pBCNF5603 | 80.67% | 0.68 |  |  |  |  |  |  |
| 96 | PH127 | FP | 0 | 0 | 0 |  |  |  |  |  |  |  |  |  |
| 97 | PH128 | FP | 0 | 1 | 10 | pCPF4969 | 96.33% | 1.78 |  |  |  |  |  |  |
| 98 | PH129 | FP | 0 | 1 | 10 | pCPF4969 | 95.20% | 1.66 |  |  |  |  |  |  |
| 99 | PH130 | FP | 0 | 1 | 10 | pCPF4969 | 95.79% | 1.64 |  |  |  |  |  |  |
| 100 | PH132 | FP | 0 | 1 | 10 | pCPF4969 | 90.80% | 1.37 |  |  |  |  |  |  |
| 101 | PH133 | FP | 0 | 1 | 10 | pCPF4969 | 94.54% | 1.58 |  |  |  |  |  |  |
| 102 | PH134 | FP | 0 | 1 | 10 | pCPF4969 | 97.08% | 1.77 |  |  |  |  |  |  |
| 103 | PH135 | FP | 0 | 0 | 0 |  |  |  |  |  |  |  |  |  |
| 104 | PH136 | FP | 0 | 0 | 0 |  |  |  |  |  |  |  |  |  |
| 105 | PH143 | FP | 0 | 1 | 10 | pCPF4969 | 96.23% | 1.48 |  |  |  |  |  |  |
| 106 | PH147 | FP | 0 | 1 | 10 | pCPF4969 | 90.89% | 0.95 |  |  |  |  |  |  |
| 107 | PH154 | FP | 0 | 1 | 10 | pCPF4969 | 97.39% | 1.42 |  |  |  |  |  |  |
| 108 | PH155 | FP | 0 | 1 | 10 | pCPF4969 | 97.50% | 1.43 |  |  |  |  |  |  |
| 109 | PH156 | FP | 0 | 0 | 0 |  |  |  |  |  |  |  |  |  |
